# Segregated basal ganglia output pathways correspond to genetically divergent neuronal subclasses

**DOI:** 10.1101/2024.08.28.610136

**Authors:** Alana I. Mendelsohn, Laudan Nikoobakht, Jay B. Bikoff, Rui M. Costa

## Abstract

The basal ganglia control multiple sensorimotor behaviors though anatomically segregated and topographically organized subcircuits with outputs to specific downstream circuits. However, it is unclear how the anatomical organization of basal ganglia output circuits relates to the molecular diversity of cell types. Here, we demonstrate that the major output nucleus of the basal ganglia, the substantia nigra pars reticulata (SNr) is comprised of transcriptomically distinct subclasses that reflect its distinct progenitor lineages. We show that these subclasses are topographically organized within SNr, project to distinct targets in the midbrain and hindbrain, and receive inputs from different striatal subregions. Finally, we show that these mouse subclasses are also identifiable in human SNr neurons, suggesting that the genetic organization of SNr is evolutionarily conserved. These findings provide a unifying logic for how the developmental specification of diverse SNr neurons relates to the anatomical organization of basal ganglia circuits controlling specialized downstream brain regions.

## INTRODUCTION

The assembly of anatomically distinct neural circuits underlies how the nervous system controls different aspects of behavior. The basal ganglia are thought to be arranged as a series of anatomically segregated and topographically organized subcircuits that modulate multiple sensorimotor, cognitive and motivated behaviors in parallel.^1^ Understanding the logic of this organization is critical for determining how dysfunction in specific basal ganglia subcircuits contributes to the particular neuropsychiatric symptomologies observed in disorders of the basal ganglia, such as movement disorders and obsessive-compulsive disorder. Identifying the genetic and molecular determinants of basal ganglia anatomical organization is also essential for developing precision therapeutics to modulate disease-relevant basal ganglia subcircuits.

Whole-brain mapping approaches have provided considerable and detailed information about the exquisite organizational logic of cortico-basal ganglia loops. Different cortical areas project to different subdomains of the striatum, the main input region of the basal ganglia.^2^ These striatal subdomains project to different subdomains of the basal ganglia’s output nuclei, the endopeduncular nucleus, globus pallidus externus (GPe) and substantia nigra pars reticulata (SNr).^3^ Different basal ganglia output subdomains in turn project to distinct targets in the thalamus, midbrain and hindbrain.^4,5^ However, the correspondence between the anatomical organization of different basal ganglia output circuits, its assembly during development, and the molecular diversity of neuronal cell types remains less understood.

We hypothesized that the anatomical organization of the SNr, the primary output nucleus of the basal ganglia, is likely to reflect different genetic lineages because the SNr is formed by distinct progenitor populations that settle in different anatomical territories. One SNr population derives from Nkx6.2^+^;Six3^+^ caudal midbrain progenitors which migrate into the anterior SNr.^6,7^ Another SNr population derives from ventral rhombomere r1, which generates Zfpm2^+^;Pax5^+^ postmitotic neuronal precursors that migrate into the ventral midbrain to form the posterior SNr.^6,8,9^ Expression of these transcription factors in adult SNr neurons appears to reflect these developmental lineages.^10,11^ Still, it remains to be determined how the diverse cell types present in SNr correspond to its anatomical organization.

Here, we use single-nucleus transcriptomics to demonstrate that the SNr is comprised of genetically distinct subclasses that appear to reflect the divergent lineages contributing to the SNr’s formation during embryonic development. We show in mouse that these genetically defined subclasses are topographically organized within SNr, project to distinct targets in the midbrain and hindbrain and receive inputs from different striatal subregions. Finally, we show that the subclasses we identified in mouse are also identifiable in the human SNr, suggesting that the genetic organization of SNr is evolutionarily conserved. Together, these findings provide a unifying logic for how the developmental specification of diverse SNr neurons relates to the anatomical organization of basal ganglia circuits controlling specialized downstream brain regions.

## RESULTS

### Identification of SNr neuronal subclasses with distinct topographic distributions

To probe the genetic diversity and topographic organization of SNr neuronal subtypes, we performed single-cell profiling of SNr neurons using single-nucleus RNA-sequencing (snRNAseq). We microdissected the SNr region from 9 12-week old mice, then performed nuclear isolation and NeuN labeling prior to fluorescence-activated nuclei sorting (FANS) to collect DAPI^+^;NeuN^+^ nuclei for 10x sequencing (Figure 1A).

**Figure 1.**
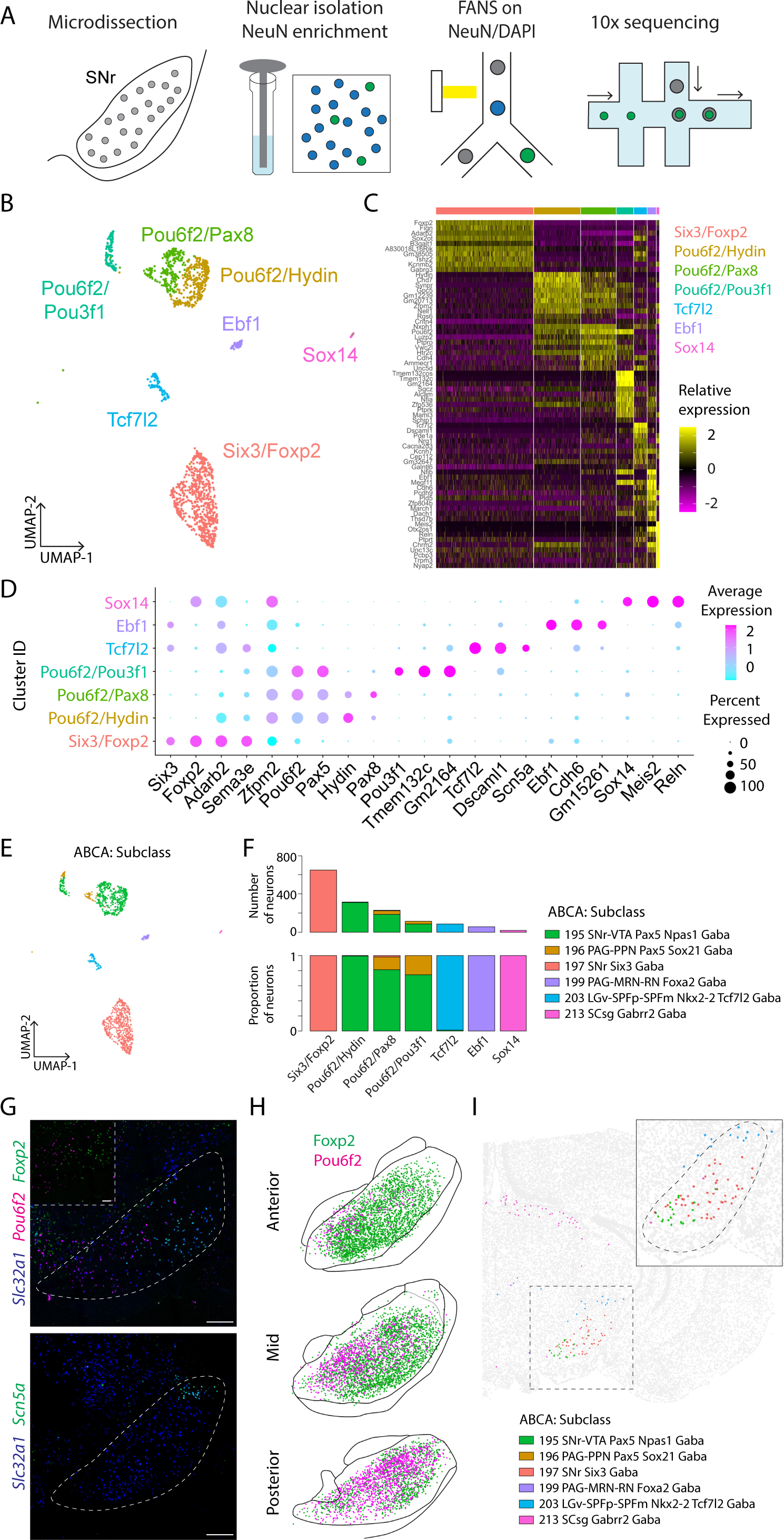
A molecular census of SNr neuronal cell types. (A) NeuN enrichment strategy for droplet-based single-nucleus RNAseq profiling of SNr neurons. (B) UMAP representation of 1499 SNr GABAergic neurons colored by assigned cluster identity. (C) Heatmap showing relative expression of cluster-enriched genes. (D) Dotplot showing expression of selected marker genes across SNr clusters. (E) SNr cluster identity mapped to Allen Brain Cell Atlas mouse taxonomy with MapMyCells. UMAP representation of SNr GABAergic neurons from 1B colored by ABCA subclass. (F) Number and proportion of neurons per cluster mapping to corresponding ABCA subclass. (G) RNAscope reveals that SNr cluster markers display different topographic distributions within SNr. Scale bars are 250µm, inset panel scalebar is 100µm. (H) Reconstruction of coordinate locations of Foxp2^+^ and Pou6f2^+^ neurons labeled by immunohistochemistry within SNr. (I) Topographic organization of ABCA subclasses from ABCA MERFISH of whole coronal mouse brain. See also Figure S1.

After exclusion of 1403 low quality samples, 2596 samples were used for subsequent analysis. Seurat was used to perform iterative unbiased clustering at multiple resolutions to determine the optimal parameters for clustering. Classification with 20 principal components and a resolution of 0.4 yielded 12 clusters, 5 of which were determined by gene marker expression to correspond to glia, dopamine neurons or glutamatergic neurons (Figure S1A). The remaining 7 clusters, comprising 1499 neurons, were GABAergic and presumed to correspond to putative SNr neurons (Figure 1B). The largest GABAergic cluster was defined by expression of *Six3*, *Foxp2, Adarb2 and Sema3e* (Figure 1D). The next 3 largest clusters shared expression of *Zfpm2*, *Pax5* and *Pou6f2*, but varied in their expression of additional markers such as *Hydin*, *Pax8*, *Pou3f1* and *Tmem132c*. The next cluster was defined by expression of *Tcf7l2* and *Scn5a*. The remaining 2 smallest clusters were defined by *Ebf1* and *Sox14* expression, respectively. We screened gene markers identified from this screen using the Allen Brain Atlas to confirm gene expression in SNr and found that all clusters featured genes expressed in SNr except the Ebf1 and Sox14 clusters, whose markers were expressed in neighboring regions.

We next determined how these cluster designations corresponded to the hierarchical taxonomy developed in the Allen Brain Cell Atlas (ABCA).^11,12^ We mapped this SNr dataset to the ABCA taxonomy using the MapMyCells function and observed that each cluster mapped almost completely to a single subclass label (Figure 1E, F). Importantly, we noted that the Six3/Foxp2 cluster mapped to one subclass, the 3 Pax5/Pou6f2 clusters mapped to the same second subclass and the Tcf7l2 cluster mapped to a third subclass. This pattern was maintained at the level of supertype, with one exception being that the Pax5/Pou6f2/Pou3f1 cluster mapped to a different supertype than the other 2 Pax5/Pou6f2 clusters (Figure S1B). Only at the ABCA cluster level did we observe assigned clusters corresponding to multiple taxonomy labels (Figure S1B). These results suggest that the SNr contains genetically distinct neuronal subtypes which correspond to 3 broader subclass designations: Six3/Foxp2, Pou6f2/Pax5 and Tcf7l2.

We determined the spatial distribution of these neuronal subtypes in the adult SNr by performing *in situ* hybridization and immunohistochemistry using select gene markers. Whereas *Foxp2* was expressed more in the dorsolateral SNr, *Pax5* and *Pou6f2* were co-expressed in the ventromedial SNr along with *Zfpm2*, which was also expressed in neighboring midbrain regions (Figure 1G and S1C). The Tcf7l2 cluster marker *Scn5a* was expressed in the most dorsolateral region of the SNr corresponding to the substantia nigra lateralis (SNL). This same topographic organization was observed after examining the spatial locations of the 3 SNr subclasses in the ABCA MERFISH dataset (Figure 1I).^11,13^ Together, these results demonstrate that SNr neuronal subclasses have distinct topographic distributions.

We noted that the two largest subclasses present in the SNr express either *Six3* or *Pax5*, suggesting a correspondence to the Six3^+^ progenitor population that forms the anterior SNr and the Zfpm2^+^;Pax5^+^ progenitor population that forms the posterior SNr. We thus asked whether the Six3/Foxp2 and Pou6f2/Pax5 subclasses are accordingly organized across the anterior-posterior axis within the adult SNr. We performed immunohistochemistry on serial coronal brain sections with antibodies against Foxp2 and Pou6f2 and found that Foxp2 distribution is biased towards the anterior SNr, whereas Pou6f2 is biased towards the posterior SNr (Figure 1H). These results suggest that the separate progenitor populations that form the SNr contribute to the spatial organization of its diverse neuronal subtypes, with the Six3/Foxp2 subclass corresponding to the Nkx6.2^+^;Six3^+^ progenitor population that forms the anterior SNr and the Pou6f2/Pax5 subclass corresponding to the Zfpm2^+^;Pax5^+^ progenitor population that forms the posterior SNr.

Lastly, we asked how these patterns of gene marker expression correspond to the known SNr marker, *Pvalb*, which is expressed in 80% of SNr neurons.^5^ We found that while the proportion of *Foxp2*^+^ neurons expressing *Pvalb* by *in situ* hybridization was higher than that of *Pou6f2^+^* and *Scn5a^+^* neurons, the difference was not statistically significant (Figure S1E). We did, however, note that *Pvalb* expressing *Foxp2*^+^ neurons express *Pvalb* at approximately 25% higher levels than *Pvalb* expressing *Pou6f2^+^* and *Scn5a^+^* neurons (Figure S1F). These results suggest *Pvalb* expression alone may not be sufficient to clearly distinguish between different SNr neuronal subclasses.

### SNr neuronal subclasses segregate by projection target

Subpopulations of SNr neurons projecting to distinct targets are also spatially organized within SNr, with neurons projecting to the superior colliculus located dorsolaterally, neurons projecting to the pons, medulla and dorsal raphe located ventromedially, and neurons projecting to the inferior colliculus located in the extreme dorsolateral subdomain (Figure S2B).^5^ We thus asked if the diverse neuronal subtypes present in the SNr corresponded to its projection targets.

To do this, we genetically profiled individual SNr neurons based on their projection to six target regions: lateral superior colliculus (LSC), medial superior colliculus (MSC), inferior colliculus (IC), dorsal raphe (DR), pontine reticular nucleus (PNo) and medullary reticular nucleus (Med). We injected AAVretro-hSyn-H2B-mCherry into one target region per mouse to retrogradely label SNr neurons that project to that region and microdissected the SNr 3 weeks later (Figure 2A). Pooling across 3-9 mice for each target region, we then isolated nuclei and performed FANS to sort mCherry^+^ nuclei into 96-well plates for snRNAseq, collecting 192-480 nuclei per target region. After exclusion of 100 low quality samples, 2012 projection-tagged samples were then merged with the unclustered 10x dataset for integration and clustering (Figure S2C). Glia and non-GABAergic clusters were excluded as before, yielding 9 GABAergic clusters, comprising 3049 samples split evenly across the 10x and projection-tagged datasets (Figure S2D). We then mapped the GABAergic clusters to the ABCA taxonomy.

**Figure 2.**
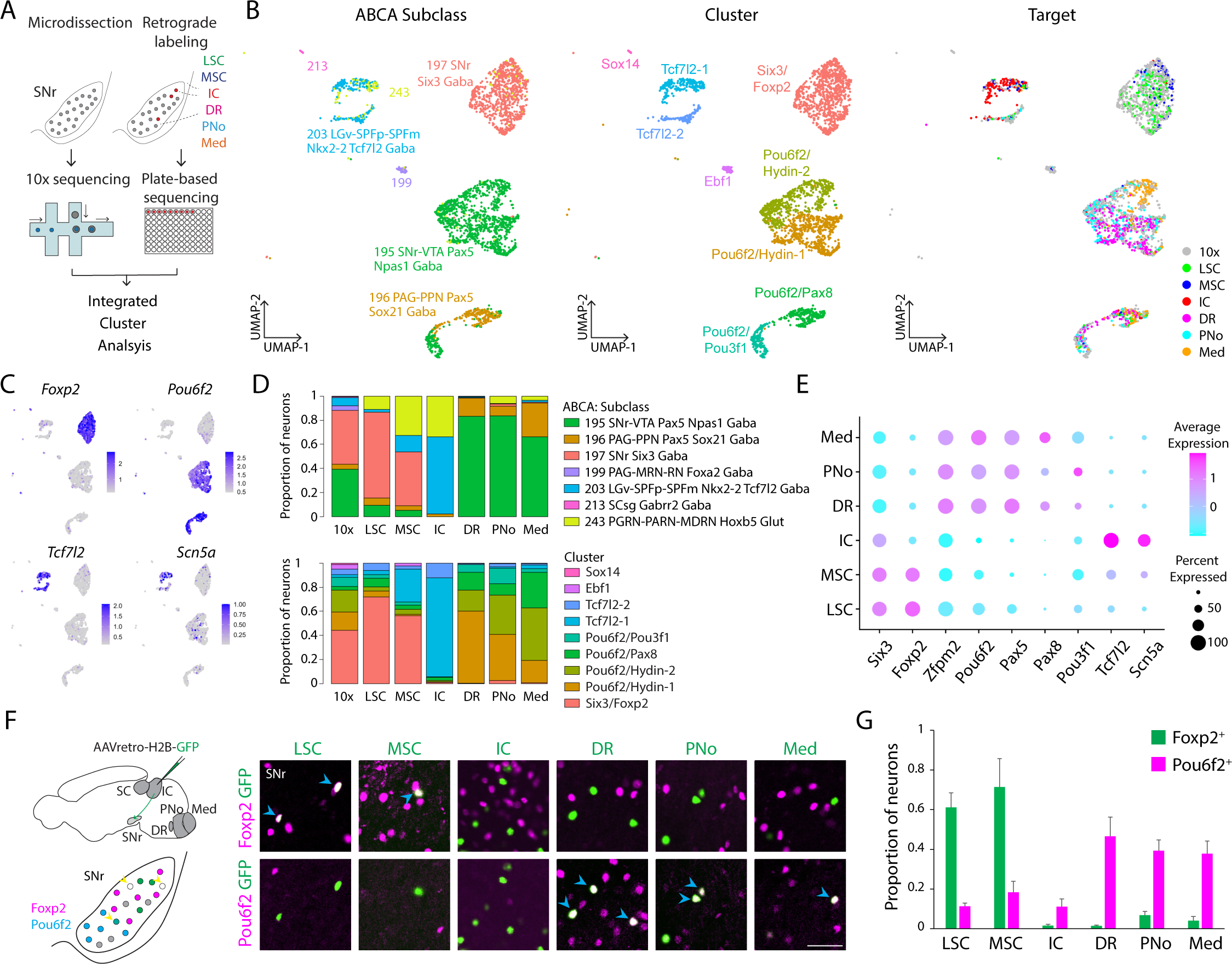
Segregation of SNr subclasses by projection target. (A) Experimental schematic for single-nucleus RNAseq profiling of SNr neurons retrogradely labeled by injection of AAVretro-hSyn-H2B-mCherry in each of six target sites. (B) UMAP representation of 3049 SNr GABAergic neurons from integrated cluster analysis of whole SNr 10x (1507 neurons) and projection-tagged plate-based RNAseq (1542 neurons), colored by mapped ABCA subclass (left), assigned cluster (middle) and projection target (right). (C) Feature plots showing single-nucleus gene expression of *Foxp2*, *Pou6f2*,*Tcf7l2* and *Scn5a*. (D) Proportion of 10x or retrogradely labeled SNr neurons corresponding to a given cluster or mapped ABCA subclass. (E) Dotplot showing expression of selected marker genes across retrogradely labeled SNr neurons. (F) Immunostaining after retrograde labeling of SNr neurons by injection of AAVretro-hSyn-H2B-GFP in each of six target sites reveals pattern of colocalization of retrogradely labeled SNr neurons with Foxp2 or Pou6f2. The same brain section is shown in the top and bottom panels for each target site. Scalebar is 50µm. (G) Quantification of (F), showing proportion of retrogradely labeled SNr neurons colocalizing with Foxp2 or Pou6f2 immunolabeling. Data are represented as mean ± SEM. See also Figure S2.

Strikingly, we observed that most SNr neurons retrogradely labeled from a given target mapped to a single ABCA subclass. 71% and 45% of SNr neurons retrogradely labeled from LSC or MSC, respectively, mapped to the Six3/Foxp2 ABCA subclass, versus 10% and 5% to the Pax5/Pou6f2 subclass and 2% and 14% to the Tcf7l2 subclass (Figure 1D). 64% of SNr neurons retrogradely labeled from IC (SNr-IC) mapped to the Tcf7l2 subclass, whereas no SNr-IC neurons mapped to the other two SNr subclasses. 83% of SNr-DR, 84% of SNr-PNo and 66% of SNr-Med neurons mapped to the Pax5/Pou6f2 subclass, with only a maximum of 2% mapping to the other two SNr subclasses. We observed a nearly identical pattern of mapping at the level of ACBA supertype, noting that the supertype corresponding to the Pou6f2/Pou3f1 cluster was evenly represented across SNr-DR, SNr-PNo and SNr-Med neurons (Figure S2G). Together these results suggest that neurons projecting to the superior colliculus, inferior colliculus and hindbrain map to different SNr subclasses. These results are further supported by our observation that the spatial distributions of SNr subpopulations projecting to different targets match the topographic distributions of their corresponding subclass type.

We next asked whether SNr neurons projecting to a single target could be distinguished with a higher level of clustering resolution. We first examined the proportional representation of the 9 GABAergic clusters from our integrated analysis across labeled SNr neurons, identifying each cluster based on its expression of previously characterized markers (Figure S2E). We found while SNr neurons projecting to DR, PNo and Med differed in the relative proportions mapping to each of the 4 Pax5/Pou6f2 clusters, there seemed to be no clusters that specifically corresponded to one SNr-hindbrain projection target versus another (Figure 2D). Similarly, we noted no patterns of ABCA cluster representation that specifically distinguished between SNr-superior colliculus neurons or SNr-hindbrain neurons (Figure S2G). These results suggest that SNr projection topography is mainly organized at the level of neuronal subclass.

We next validated the subclass correspondence of projection-defined SNr subtypes *in vivo* by combining retrograde labeling with histologic marker expression. We found that ∼60-70% of SNr neurons retrogradely labeled from LSC or MSC expressed Foxp2, whereas only 1-7% of SNr neurons retrogradely labeled from IC or hindbrain expressed Foxp2 (Figure 2F and 2G). In contrast, we found that 38-47% of SNr neurons retrogradely labeled from DR, PNo or Med expressed Pou6f2, whereas only 11-18% of SNr neurons retrogradely labeled from LSC, MSC or IC expressed Pou6f2. Less than 11% of SNr-IC neurons expressed either Foxp2 or Pou6f2. Instead, SNr-IC neurons colocalized with expression of *Scn5a* by *in situ* hybridization (Figure S2H). Taken together, these results demonstrate that SNr neurons projecting to the superior colliculus, inferior colliculus and hindbrain map to different SNr subclasses expressing different gene markers.

### SNr neuronal subclasses have distinct brain-wide targets

We next sought to characterize the brain-wide projection patterns of 3 SNr subclasses. To do this, we used genetic and anatomical strategies to access these subclass populations with viral labeling. To access the SNr Six3/Foxp2 subclass, we injected AAV-Con/Fon-syn-Myc into the SNr of Foxp2-Cre;Vgat-Flp mice, restricting expression to GABAeric Foxp2^+^ neurons in the dorsolateral SNr (Figure 3A and S3B). To access the SNr Pax5/Pou6f2 subclass, we first validated use of a Pou6f2-Flp mouse by *in situ* hybridization, showing that 70% of endogenous *Pou6f2*^+^ SNr neurons are labeled after crossing Pou6f2-Flp to a reporter mouse line (Figure S3C and S3D). As expected, injection of AAV-FRT-mGFP-synaptophysin-mRuby into the SNr of Pou6f2-Flp mice labeled neurons in the ventromedial SNr (Figure 3A). Lastly, given that the Tcf7l2 subclass was restricted within SNr to SNr-IC neurons, we accessed this population with an intersectional anatomical strategy, injecting AAVretro-Flpo into the IC and a AAV-FRT-mGFP-synaptophysin-mRuby into the dorsal SNr. This strategy labeled neurons in the extreme dorsolateral subdomain of SNr (Figure 3A).

**Figure 3.**
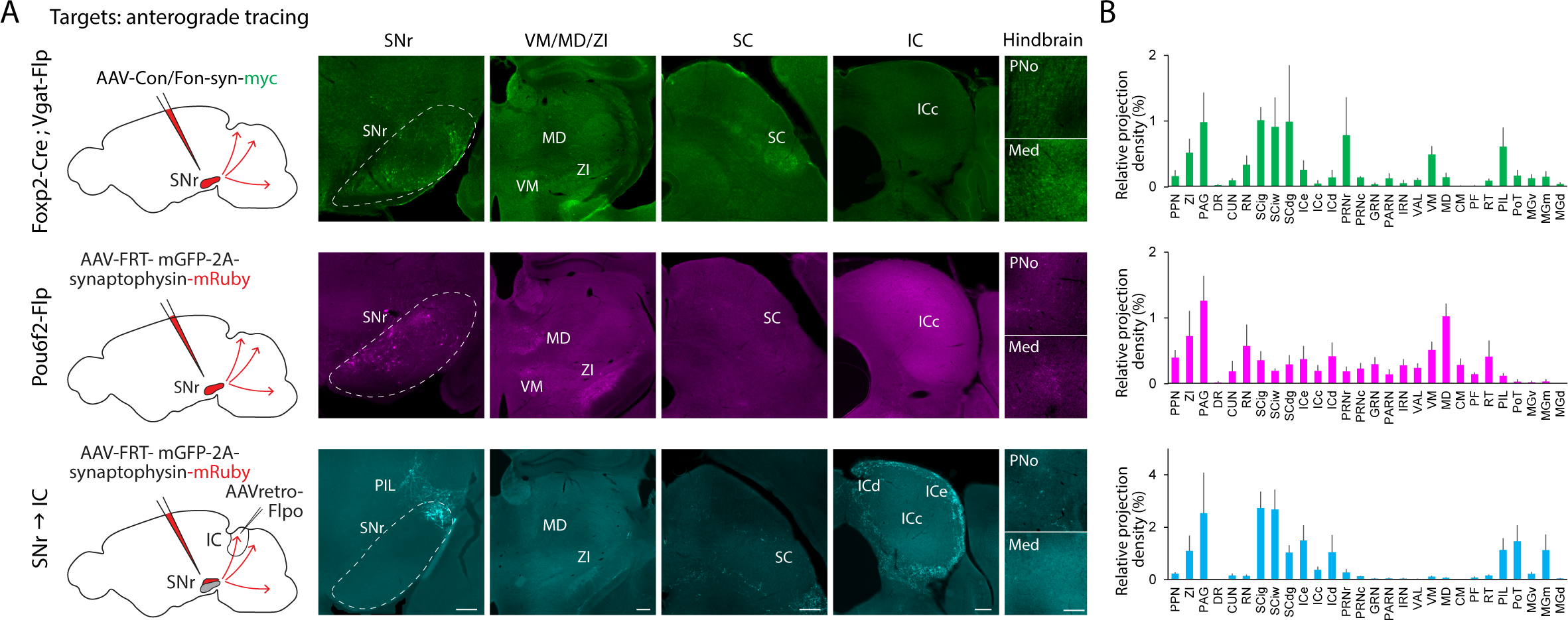
SNr subclasses have different projection targets. (A) Experimental schematic and representative images for brain-wide anterograde tracing from three SNr subclasses. Scale bars are 250µm. (B) Quantification of relative axon density across brain regions (see Table S1 for brain region acronyms). N = 3 mice for Foxp2 and Pou6f2 subclasses and 4 mice for SNL-IC subclass. Data are represented as mean ± SEM. See also Figure S3.

Consistent with previous work, we found that all SNr subclasses sent projections to known common targets such as pedunculopontine nucleus, zona incerta and periaqueductal grey.^5^ However, we noted that each population differed in its projections to specialized downstream targets (Figure 3A and 3B). As expected, the Foxp2 subpopulation mainly projected to the superior colliculus, whereas the Pou6f2 subpopulation sent projections to the pons and medulla and the SNr-IC subpopulation projected to the shell of the inferior colliculus, with additional collaterals to the superior colliculus. The 3 SNr subclasses also differed in their projections to thalamic targets. The Foxp2 subpopulation sent its densest thalamic projection to the ventral medial nucleus of the thalamus, whereas the Pou6f2 subpopulation projected to the mediodorsal nucleus of thalamus and the SNr-IC subpopulation projected to the posterior intralaminar thalamic nucleus, posterior triangular thalamic nucleus and medial geniculate complex. Together, these results demonstrate that genetically distinct and topographically segregated SNr subclasses have distinct brain-wide projections.

### SNr neuronal subclasses receive input from different striatum subdomains

Because SNr subclasses occupy distinct spatial subdomains within SNr, we hypothesized that they would also receive inputs from different subdomains within the striatum. To test this, we sought to characterize the spatial distribution of striatal inputs to each SNr subclass using presynaptic modified rabies tracing. We injected rabies helper viruses using the following strategy. We delivered rabies helper viruses to the SNr Six3/Foxp2 subclass by injecting a 1:1 mixture of AAV-Con/Fon-TVA-mCherry and AAV-Con/Fon-N2cG into the SNr of Foxp2-Cre;Vgat-Flp mice. We accessed the Pou6f2 subclass by injecting a 1:1 mixture of AAV-FlpX-TVA-EYFP and AAV-FlpX-mKate-N2cG into the SNr of Pou6f2-Flp mice. Lastly, we accessed the SNr-IC subclass by injecting AAVretro-Flpo into the IC and 1:1 mixture of AAV-FlpX-TVA-EYFP and AAV-FlpX-mKate-N2cG into the dorsal SNr. 3 weeks after helper virus injection, we injected EnVA-N2cΔG-GFP into the SNr.

Injection of helper virus followed by rabies virus labeled both starter cells in SNr in the expected spatial location for each subclass, as well as presynaptic input cells located in the striatum (Figure 4A and S4A). As a control, we injected EnVA-N2cΔG-GFP after injection of only one helper virus. As expected, rabies injection after sole injection of N2cG helper virus failed to label starter cells (Figure S4B), whereas rabies injection after sole injection of TVA helper virus labeled starter cells in SNr but failed to label monosynaptic input cells in the striatum (Figure S4C).

**Figure 4.**
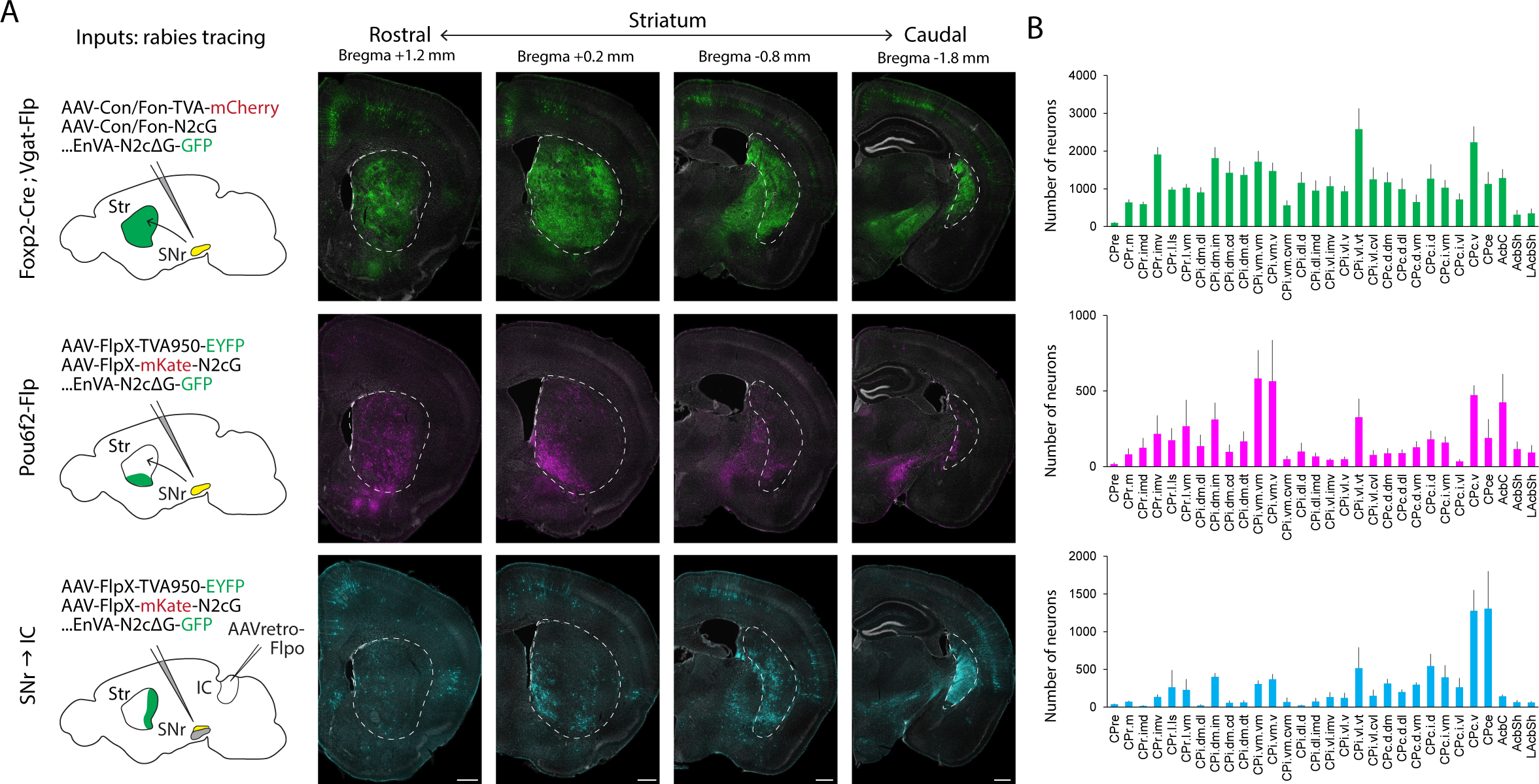
SNr subclasses receive input from different striatum subdomains. (A) Experimental schematic and representative images for rabies virus presynaptic input tracing from three SNr subpopulations. Scale bars are 500 (B) Quantification if number of neurons across striatum subdomains (see Table S2 for striatum subdomain acronyms). N = 4 mice for Foxp2 subclass, 3 mice for Pou6f2 subclass and 3 mice for SNL-IC subclass. Data are represented as mean ± SEM. See also Figure S4.

After aligning brain sections from the rabies experiments to a common coordinate framework, we next determined the number of presynaptic input cells across all striatum subdomains (Figure 4B). For the Foxp2 SNr subclass, we observed labeled presynaptic input cells across most striatum subdomains. In contrast, we found that presynaptic inputs to the Pou6f2 SNr subclass were biased towards ventromedial subdomains of the striatum and that presynaptic inputs to the SNr-IC subclass were biased towards the tail of the striatum. These results show that genetically distinct SNr subclasses receive overlapping, but non-identical patterns of input from the striatum.

### SNr neuronal subclasses are conserved in human

Recent work has suggested that the neural cell types and patterns of gene expression in the midbrain are largely conserved across mouse and human.^14^ We thus wondered with the neural subclasses we identified in mouse are also present in the human brain. To test this, we examined a published dataset of human midbrain neurons profiled with snRNAseq and grossly annotated by cell type.^15^ We used Harmony and Seurat to integrate and cluster 39110 human non-dopaminergic neurons from 8 control donors. Clustering with 20 principle components and a resolution of 0.4 yielded 23 clusters. After excluding 4 clusters of low-quality samples and 11 glutamatergic clusters, we identified 8 remaining clusters, comprising 9122 samples, corresponding to GABAergic neurons (Figure S5A).

We screened these human GABAergic clusters for markers previously identified in mouse and found that 6 of the 8 clusters expressed known markers in the same configurations observed in mouse (Figure 5). 2 human GABAergic clusters co-expressed *SIX3*, *FOXP2*, *ADARB2* and *SEMA3E*, analogous to the Six3/Foxp2 SNr subclass identified in mouse. These clusters differed primarily in their relative expression *PVALB*. 2 clusters co-expressed *ZFPM2*, *POU6F2* and *PAX5*, analogous to the Pax5/Pou6f2 SNr mouse subclass, with one cluster expressing higher levels of *PAX8* and the other expressing higher levels of *TMEM132C*. 1 cluster expressed *TCF7L2*. The last cluster co-expressed *FOXP2*, *ZFPM2*, *MEIS2*, *OTX2* and *RELN*, analogous to the Sox14 cluster in mouse which appeared not to correspond to SNr. These observations suggest a remarkable correspondence between the neural subtypes and gene expression patterns of the SNr in mouse and human.

**Figure 5.**
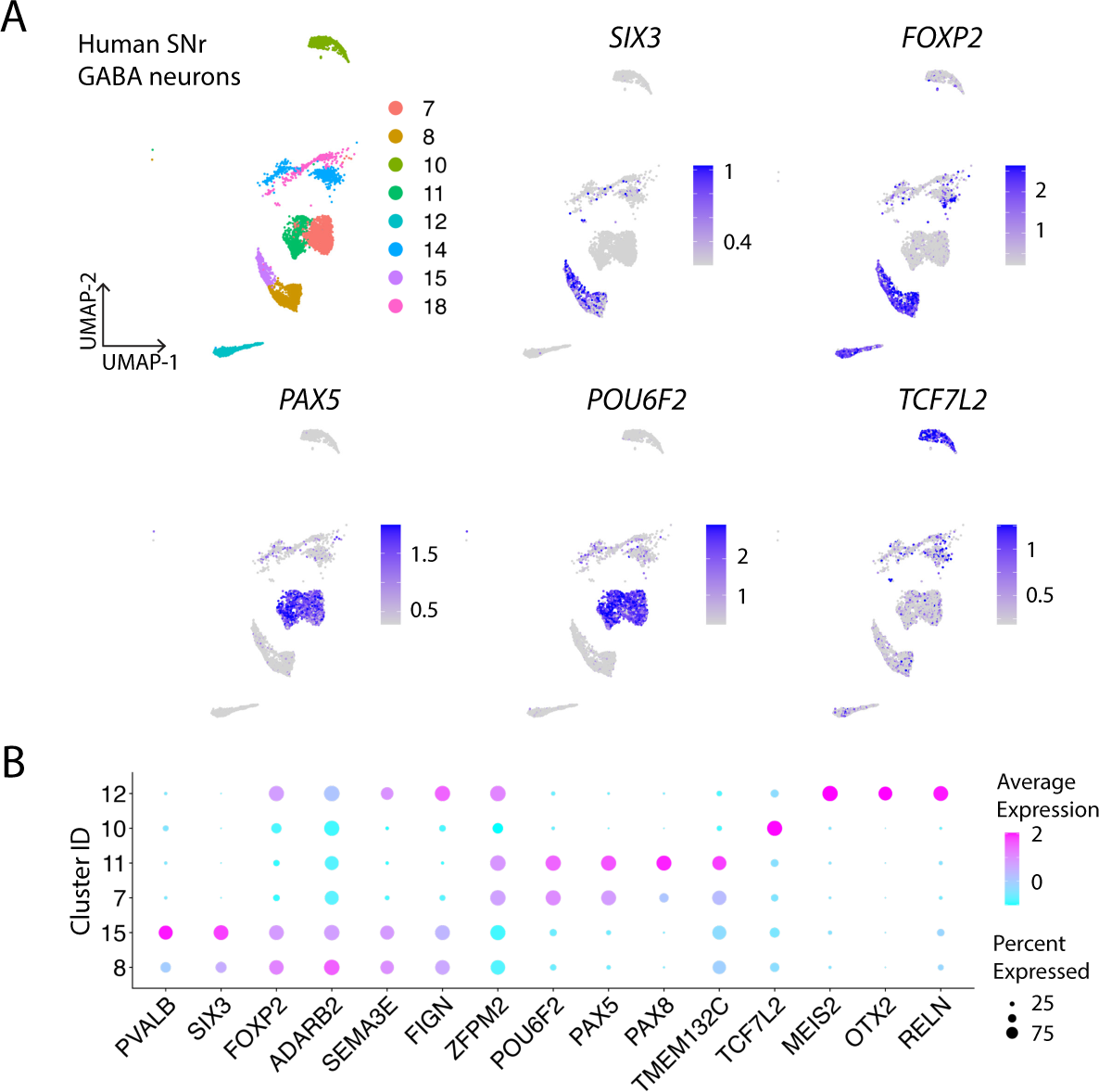
Conserved subclass marker gene expression in human SNr. (A) UMAP representation of 9122 human GABAergic SNr neurons identified after clustering 39110 non-dopaminergic midbrain neurons from 8 human donors. Feature plots showing relative expression of SNr markers. (B) Dotplot showing expression of selected marker genes across human SNr clusters. See also Figure S5.

### Leveraging neural subclasses to interrogate disease pathology

Characterizing cell type diversity is an important prerequisite for determining cell type-specific patterns of gene dysregulation observed in disease states. Recent work has shown that dopamine depletion, a key component of Parkinson’s Disease, results in variable network dysfunction within SNr.^16^ We thus asked whether Parkinson’s Disease alters transcriptional dynamics in SNr neurons and whether such gene dysregulation varies across SNr cell types. To do this, we leveraged our characterization of genetically distinct, conserved neural subtypes in SNr to probe cell-type specific responses to Parkinson’s Disease (PD) in SNr. We merged 39110 human non-dopaminergic samples from 8 control donors with 34779 non-dopaminergic samples from 7 Parkinson’s Disease donors, integrating across disease state using Harmony. After performing clustering, we identified 6 GABAergic clusters comprising 13910 samples, 9307 from control donors and 4603 from Parkinson’s Disease donors (Figure S5B). Of these, 3532 samples corresponded to the SIX3/FOXP2 subtype and 6638 samples corresponded to the PAX5/POU6F2 subtype, each with similar proportional representations from control and PD donors (Figure S5C and S5D).

We next performed differential gene expression analysis between control and PD donor samples within the two largest subtypes, SIX3/FOXP2 and PAX5/POU6F2.We found that gene expression was largely preserved across these SNr subtypes between control and PD samples (Figure S5E) (Pearson correlation coefficient 0.98 for SIX3/FOXP2 and 0.99 for PAX5/POU6F2). However, the genes that were selectively upregulated in PD samples differed between the SIX3/FOXP2 and PAX5/POU6F2 subtypes (Figure S5F). These results suggest that despite massive loss of neighboring SNc dopamine neurons, gene expression in SNr is largely preserved in PD, although it may exhibit subtle differences across subclasses.

## DISCUSSION

This study shows that the detailed anatomical architecture of parallel basal ganglia loops exists within a broader molecularly organized framework comprised of genetically distinct basal ganglia output subclasses with different circuit connectivity. These genetically distinct SNr subclasses appear to correspond to the divergent progenitor populations that contribute to the formation of SNr during embryonic development and project to target regions with different sensorimotor functions, raising important questions about the evolutionary origins of this organizational scheme and its relevance to motor control. Finally, we show that SNr subclasses appear to be conserved in the human brain and introduce how they can be leveraged to probe cell type-specific dysfunction in disease states. Together, this work provides a unifying logic for how the developmental specification of diverse SNr neurons relates to the anatomical organization of basal ganglia circuits controlling specialized downstream brain regions, laying a critical foundation for future work dissecting how molecularly segregated basal ganglia output pathways differentially contribute to sensorimotor behavior and disease states.

### Conserved neuronal diversity in the SNr

Neuronal cell types can be classified according to multiple genetic, molecular, morphologic, anatomical and electrophysiologic parameters, raising considerable debate as to what level of resolution is appropriate for defining biologically meaningful neuronal cell types.^17^ It has consequently been argued that cell type designations must be considered in the context of their developmental and evolutionary specification. Previous groups have examined SNr diversity through single cell transcriptomics, anatomical tracing and electrophysiologic profiling, finding that cells in the SNr are molecular diverse and exhibit important differences in their projection patterns and physiologic properties.^5,18^ We extend this work by characterizing SNr transcriptomic diversity using a hierarchical taxonomy framework within the context of the SNr’s developmental origins.

We find that the SNr is organized at the level of subclasses, with subclasses corresponding to different developmental progenitor populations, settling in different spatial subregions of the SNr and projecting to different target regions (Figures 1 and 2). While further transcriptional differences can be detected within these subclasses by clustering SNr neurons at higher levels of resolution, SNr supertype or cluster designations do not appear to display significant differences in settling position or connectivity that would suggest that they constitute meaningfully different cell types. Instead, transcriptomic variation within each subclass may contribute to other cellular properties relevant for neuronal function. SNr neurons projecting to different hindbrain target regions, for example, correspond to the same transcriptomic subclass, but vary in their relative composition of different transcriptomic clusters and display distinct electrophysiologic signatures (Figure 2).^5^

Further support for our argument that SNr subclasses constitute meaningfully distinct cell types comes from our observation that these subclass designations appear to be conserved across species. Transcriptomic analysis of human SNr reveals the same patterns of transcription factor expression found in mouse, suggesting that human SNr exhibits the same subclass distinctions (Figure 5). Ongoing efforts to catalogue and integrate brain cell types across species will be critical to resolve to what extent the cellular diversity of SNr in rodent is maintained, or perhaps extended, in human, and to determine at what point in vertebrate evolution this diversity was generated. While the central architecture of cortico-basal ganglia loops appears to be present in early vertebrates, it is unclear how discrete modules within this architectural framework have evolved to support varied motor programs.^19^ It is tantalizing to consider that the evolutionary generation of diverse basal ganglia output cell types occurred to meet new demands in locomotor and orienting behavior.

### Developmental origins of SNr cell types

A key finding of our work is that the SNr’s transcriptomic subclass organization corresponds to the distinct progenitor pools that form the SNr in embryonic development. The SNr is formed by at least two progenitor populations, a Six3^+^ midbrain progenitor population that forms the anterior SNr and a Zfpm2^+^;Pax5^+^ ventral rhomomere r1 progenitor population that forms the posterior SNr.^6–9^ The Six3/Foxp2 and Pax5/Pou6f2 subclasses appear to derive from these two progenitor populations based on their shared transcription factor expression and settling positions in the anterior and posterior SNr, respectively. The origins of the third Tcf7l2 subclass are less clear, however. In mouse, this subclass expresses Six3, suggesting that it also derives from the Six3^+^ midbrain population. Future developmental sequencing and lineage tracing studies will be required to clarify the developmental origins of SNr cellular diversity.

An intriguing observation is that the different progenitor origins of SNr neurons appear to correspond to their differing adult connectivity. The Six3^+^ midbrain progenitor population appears to give rise to Six3^+^ adult SNr neurons that preferentially project to targets in the midbrain, whereas the Zfpm2^+^;Pax5^+^ rhombomere progenitor population appears to give rise to Zfpm2^+^/Pax5^+^/Pou6f2^+^ adult SNr neurons that preferentially project to targets in the hindbrain. How the developmental specification of these neuronal populations informs their choice of settling position and pattern of axon branching remains to be determined.

### Spatial organization and circuitry

The developmental specification of diverse subclasses in SNr informs their different settling positions within the SNr and anatomical circuitry. The Six3/Foxp2 SNr subclass settles in more anterior and dorsolateral regions of SNr, receives broad striatal inputs and sends projections to the superior colliculus. The Pax/Pou6f2 SNr subclass settles in more posterior and ventromedial regions of SNr, receives biased input from ventromedial striatum and projects to hindbrain targets. The Tcf7l2 SNr subclass occupies the most dorsolateral subdomain of SNr, receives biased input from the tail of the striatum, and projects to the shell of the inferior colliculus with collaterals to the superior colliculus. These findings are consistent with prior anatomical tracing studies examining circuit pathways linking specific striatal subdomains, SNr subregions and the downstream targets.^3–5^

These subclass designations, however, do not explain the full complexity of SNr anatomical circuity. Within each subclass, SNr neurons can be further classified based on their specific settling positions, striatal inputs and projections to specific downstream targets. Foxp2^+^ SNr neurons, for example, receive inputs across the striatum and send axon collaterals throughout the superior colliculus (Figures 3 and 4). Within this subclass, SNr neurons projecting to the central superior colliculus occupy a ventrolateral territory of SNr that receives biased input from the dorsolateral striatum, whereas SNr neurons projecting to the lateral superior colliculus occupy more dorsal regions of the SNr that receive biased input from ventrolateral striatum.^3–5^ This suggests an organizational framework in which developmental genetic programs establish the broad genetic, topographic and anatomical distinctions between subclasses, upon which further axon guidance and molecular programs guide the formation anatomically and electrophysiologically specialized circuitry.

### Implications for behavior and disease

Basal ganglia output circuits are thought to modulate behavior through a combination of specialized projections that engage discrete circuit modules and broad collateralization to more generalized motor output targets.^1^ Our work reinforces this view and further suggests that the molecular segregation of SNr subclasses supports engagement of categorically distinct sensorimotor functions. Our work would suggest that Six3/Foxp2 SNr neurons are more involved in orienting behaviors through their projections to the superior colliculus, that Pax5/Pou6f2 SNr neurons are more involved in regulating locomotion through their projections to the hindbrain and that Tcf7l2 SNr neurons regulate sensory-driven motor behaviors through their inputs from the tail of the striatum and projections to the inferior colliculus.^20–22^ The Pax5/Pou6f2 subpopulation, through its striatal inputs from ventromedial striatum and the nucleus accumbens and output to mediodorsal thalamus, may also play a specialized limbic role in reward processing.^23^ Within this overarching framework, the engagement of specific striatal subdomains and SNr subset modules may direct more specialized behaviors, such as directional behavior or orofacial behavior.^4^

Basal ganglia activity is dysregulated in multiple movement and psychiatric disorders but has been difficult to decipher given the functional heterogeneity of these circuits. Dissecting the role of anatomically or molecularly defined basal ganglia subcircuits will prove critical for linking how disease processes give rise to circuit dysfunction and specific constellations of symptoms. In Parkinson’s Disease, our work suggests that SNr circuits exhibit only subtle changes in gene expression (Figure 5). Recent work suggests, however, that SNr neuronal subsets display different electrophysiologic changes in response to dopamine loss that may explain a differential role in PD pathophysiology.^24^ Given that SNr is a current target of deep brain stimulation therapy,^25^ clarifying the regional and molecular specialization of SNr subsets may prove useful for guiding and developing more precise, cell-type specific therapeutic approaches.

## ACKNOWLEDGEMENTS

We thank Ira Shieren and Zuckerman Institute’s Flow Cytometry platform for assistance with FACS. We thank Erin Bush for providing guidance on the design of the RNA sequencing experiments and Zack Lewis for feedback on RNA sequencing analysis. We thank Susan Brenner-Morton for antibodies. We are grateful to David Ng for producing custom recombinant DNA plasmids with strategic input from Akira Fushiki. We thank Katie Balaguer for producing exceptional viruses. We thank Luke Hammond and Humberto Ibarra for assistance with imaging and analysis. We thank Ed Callaway (Salk) for providing the AAVretro-H2B-GFP plasmid and Silvia Arber (FMI) for providing the AAV-DJ-EF1a-ConFon-Synaptophysin-10Xmyc-WPRE plasmid. Single cell sequencing was performed with support from the Columbia Genome Center, which is funded in part through the NIH/NCI Cancer Center Support Grant P30CA013696 and NIH National Center for Advancing Translational Sciences Grant UL1TR001873. CVS-N2c rabies viruses were produced by the Center for Neuroanatomy with Neurotropic Viruses, supported by P40 OD010996. Imaging was performed with support from the Zuckerman Institute’s Cellular Imaging platform. This work was supported by an NIH T32 award (T32MH015144), a Leon Levy Fellowship in Neuroscience and a BBRF Young Investigator Award to A.M. and National Institute of Health funding (5U19NS104649) and the Aligning Science Across Parkinson’s (ASAP-020551) through the Michael J. Fox Foundation for Parkinson’s Research (MJFF) to R.M.C. The authors have applied a CC BY NC public copyright license for the manuscript.

## AUTHOR CONTRIBUTIONS

A.M. designed and conducted the experiments, performed the analysis, discussed the results and wrote the paper. L.N. performed histology and imaging analysis. J.B. provided the transgenic Pou6f2-Flp mouse line. R.M.C. designed the experiments and analyses, discussed the results, and edited the paper.

## MATERIALS AND METHODS

### Experimental model and animals

All experiments and procedures were performed according to National Institutes of Health (NIH) guidelines and approved by the Institutional Animal Care and Use Committee of Columbia University. Adult mice aged 2–6 months, were used for all experiments. The strains used were: C57BL6/J (Jackson Laboratories, 000664), Foxp2-Cre (Jackson Laboratories, 030541), Vgat-Flp (Jackson Laboratories, 029591), Pou6f2-Flp (gift from Jay Bikoff) and Ai65F (Jackson Laboratories, 032864). All mice were kept under a 12-h light–dark cycle.

### Pou6f2-Flp mouse generation

The *Pou6f2::Flpo* line was generated using CRISPR/Cas9 technology to insert a P2A-Flpo cassette directly before the stop codon in Exon 8. Briefly, single guide RNAs (sgRNAs; Synthego) targeting within 20bp of the desired integration were designed with at least 3bp of mismatch between the target site and any other site in the genome. Prior to embryo injection, sgRNAs were tested for activity in mouse Neuro2a cells stably expressing Cas9 and assayed by targeted next-generation sequencing (NGS). For animal model generation, 3 to 4 week old C57BL/6J female mice were superovulated with 5 units of pregnant mare’s serum gonadotropin (ProSpec), followed 48 hours later with 5 units of human chorionic gonadotrophin (Sigma). After overnight mating with C57Bl/6J males, females were euthanized and oocytes were harvested from the ampullae. The protective cumulus cells were removed with hyaluronidase, oocytes were washed, and fertilized zygotes were injected into the pronucleus with 10-20 ng/ml of sgRNA, 30-60 ng/ml SpCas9 protein (St. Jude Protein Production Core), and 5 ng/ml of ssDNA donor (megamer, IDT). Following injection, oocytes were returned to culture media (M16 or Advanced-KSOM, Millipore) and transferred to Day 0.5 pseudopregnant foster mice. Founder mice were genotyped by targeted next-generation sequencing followed by analysis using CRIS.py, and validated by immunohistochemistry to confirm recombination in Pou6f2-expressing neurons.

### Molecular cloning and recombinant AAV production

AAV transfer plasmids were generated through modifications of existing plasmids using HIFI DNA assembly cloning (NEB). pAAV-Ef1a-Con/Fon N2cG, based on pAAV-Ef1a-Con/Fon oG (Addgene #131778), was made by replacing coding sequences from the rabies oG protein with orthologous sequences from N2cG (PCR amplified from Addgene #73481). High scoring predicted splice donor and splice acceptor sites on the N2cG sense sequence were mitigated by introduction of synonymous codon substitutions to enhance protein expression.^26^ pAAV-ehSyn-fDIO-TVA950-eYFP, based on pAAV-DIO-TVA950-eYFP (Addgene #120269), was generated by replacement of the DIO cassette with a fDIO cassette (synthesized by GenScript), and insertion of the TVA950-eYFP sequence. pAAV-CAG-fDIO-mKate2-2a-N2cG was generated by HIFI DNA assembly of mKate2 (from Addgene #159928), T2A and N2cG ORFs (from Addgene #73481) into a pAM-FLEX AAV2 backbone that had been modified to replace DIO with fDIO sequences.^27^ PCR amplicons were generated using Q5 DNA polymerase (NEB). Plasmids were propagated at 30°C in Stable cells (NEB). To produce recombinant AAVs, HEK293T cells (ATCC) were triple transfected using polyethylenimine (PEI) as previously described.^28^ AAVs aliquots were stored at -80C until experimental use.

### Stereotaxic viral injections

All surgeries were performed under sterile conditions and isoflurane (1-3%) was used to anaesthetize mice. Throughout each surgery, mouse body temperature was maintained at 37° C using an animal temperature controller (ATC2000, World Precision Instruments). Analgesia in the form of subcutaneous injection of carprofen (5 mg per kg body weight) or buprenorphine XR (0.5– 1 mg per kg body weight) was administered on the day of the surgery, along with bupivacaine (2 mg per kg body weight). Mice were placed in a stereotaxic holder (Kopf) and a midline incision was made to expose the skull. A craniotomy was made over the injection site with a drill (Osada, Los Angeles, CA, USA). All injections were performed using a Nanoject II Injector (Drummond Scientific, Broomall, PA, USA) at a rate of 4.6 nl every 5 s. Pulled glass injection pipettes were left in place for 5 min after injection to allow for virus absorption, and incisions were closed with Vetbond tissue adhesive (3M, Maplewood, MN, USA). Mice were subsequently allowed to recover from the anesthesia in their homecage on a heating pad.

To retrogradely label SNr neurons based on their projection targets, AAVretro-hSyn-H2B-mCherry (Salk Institute, 1.32 x 10^13^ GC/ml) or AAVretro-hSyn-H2B-GFP (ZI Virology, 1.10 x 10^13^ GC/ml) were injected into one of six hindbrain regions: lateral superior colliculus (75-125 nl, AP: -3.6 mm, ML: 1.3 mm, DV: 2.1 mm below brain surface), medial superior colliculus (125-150 nl, AP: -3.6 mm, ML: 0.35 mm, DV: 1.3 mm), inferior colliculus (75-125 nl, AP: -5 mm, ML: 1.3 mm, DV: 1.3 mm), dorsal raphe (150 nl, AP: -4.5 mm, ML: 1.12 mm, DV: 3.1 mm at 22°), pontine reticular nucleus (100-125 nl, AP: -4.5 mm, ML: 0.8 mm, DV: 4.7 mm), medullary reticular nucleus (200 nl, AP: -6.4 mm, ML: 1.2 mm, DV: 4.8 mm). Mice were sacrificed 3 weeks later.

To label SNr neurons in Foxp2-Cre;Vgat-Flp mice, 400 nl of AAV(DJ)-hSyn-Con/Fon EYFP-WPRE (UNC Vector Core, 5.40 x 10^12^ GC/ml) was injected into SNr (AP: -3.2 mm, ML: 1.4 mm, DV: 4.6 mm). To anterogradely label SNr neurons in Foxp2-Cre;Vgat-Flp mice, 200 nl of AAV-DJ-EF1a-ConFon-Synaptophysin-10Xmyc-WPRE (ZI Virology, 5.40 x 10^12^ GC/ml) was injected into the right SNr. To anterogradely label SNr neurons in Pou6f2-Flp mice, 200 nl of AAV-DJ-hSyn-FLEX_FRT-mGFP-2A-synaptophysin.mRuby (Stanford Vector Core, 1.10 x 10^13^ GC/ml) was injected into right SNr. To anterogradely label SNL à IC neurons in Bl6 mice, 50 nl of AAVretro-EF1a-Flpo (Addgene 55637-AAVrg, 1.10E+13, diluted 50%) was injected into right IC and 75 nl of AAV-DJ-hSyn-FLEX_FRT-mGFP-2A-synaptophysin.mRuby (Stanford Vector Core, 1.10 x 10^13^ GC/ml, diluted 50%) was injected into right SNL (AP: -3.15 mm, ML: 1.9 mm, DV: 3.7 mm). Mice were sacrificed 4-6 weeks later.

To label presynaptic inputs to SNr neurons in Foxp2-Cre;Vgat-Flp mice, 400 nl of a 1:1 mixture of AAV8-Con/Fon-TVA-mCherry (Stanford Vector core, 1.10 x 10^13^ GC/ml, diluted 1:10) and AAV8-Con/Fon-N2cG (ZI Virology, 4.80 x 10^12^ GC/ml, diluted 50%) was injected into right SNr. 3 weeks later, 150 nl of EnVA-N2cΔG-GFP (Thomas Jefferson) was injected into right SNr. To label presynaptic inputs to SNr neurons in Pou6f2-Flp mice, 400 nl of a 1:1 mixture of pAAVDJ-Esyn-FlpX-TVA950-EYFP-WPRE (ZI Virology, 2.2 x 10^13^ GC/ml, diluted 1:10) and AAVDJ-CAG-FlpX-mKate2.0-N2cG (ZI Virology, 2.6 x 10^12^ GC/ml) was injected into right SNr. 3 weeks later, 150 nl of EnVA-N2cΔG-GFP was injected into right SNr. To label presynaptic inputs to SNL à IC neurons in Bl6 mice, 100 nl of AAVretro-EF1a-Flpo (as above, diluted 50%) was injected into right IC and 300 nl of a 1:1 mixture of pAAVDJ-Esyn-FlpX-TVA950-EYFP-WPRE and AAVDJ-CAG-FlpX-mKate2.0-N2cG (as above) was injected into right SNL. 3 weeks later, 150 nl of EnVA-N2cΔG-GFP was injected into right SNL. Mice were sacrificed 1 week after rabies injection. For rabies control experiments, the same equivalent volume of each helper virus component was used. In several control experiments, EnVA-N2cΔG-tdTomato was substituted for EnVA-N2cΔG-GFP at the equivalent volume.

### Histology

Mice were deeply anesthetized with isoflurane and transcardially perfused with PBS followed by ice-cold 4% paraformaldehyde. Brains were postfixed overnight in 4% paraformaldehyde, and then cryopreserved in a 30% sucrose solution for 3 days at 4°C. Brains were embedded in Optimum Cutting Temperature Compound (Tissue-Tek), frozen and stored at -80°C. 50μm coronal sections were cut on a cryostat. Primary antibodies used in this study were Rabbit anti-Foxp2 (Abcam, 1:10k), Guinea Pig anti-Pou6f2 (Columbia, 1:2k), Rabbit anti-GFP-488 (Invitrogen, 1:1k), Chicken anti-GFP (Aves labs, 1:2k), Rabbit anti-RFP (Rockland, 1:2k), Rat anti-mCherry (Invitrogen, 1:2k), Mouse anti-TH (Immunostar, 1:10k) and Chicken anti-Myc (Invitrogen, 1:250). Primary antibodies were diluted in PBST 0.3%, 0.5% BSA and 0.05% thimersol. Tissue sections were incubated with primary antibodies for 2 days at 4°C, washed 3 times in PBS and incubated with secondary antibodies at a 1:1000 dilution for 2 hours at room temperature. Sections were then washed 3 times in PBS and counterstained at room temperature with DAPI at a 1:1000 dilution for 15 minutes.

For fluorescence *in situ* hybridizations, mice were deeply anesthetized and transcardially perfused with PBS. Brains were rapidly dissected, frozen on dry ice in OCT, and stored at -80° C. 15-μm coronal sections were cut on a crysostat, adhered to Super-Frost Plus Slides, and immediately stored at -80° C. Sections were fixed in 4% PFA in PBS and stained according to Advanced Cell Diagonstics (ACD) RNAscope Multiplex Fluorescent V2 Assay manual. Probes were visualized using Opal dyes (Akoya Biosciences). Slides were counterstained with DAPI.

### Imaging

Automated high-throughput imaging of tissue sections was performed on a custom-built automated slide scanner using a AZ100 microscope equipped with a 4x 0.4NA Plan Apo objective (Nikon Instruments Inc) and P200 slide loader (Prior Scientific), controlled by NIS-Elements using custom acquisition scripts (Nikon Instruments Inc.). Confocal imaging was performed on a W1-Yokogawa inverted spinning disk confocal using a 10x or 20x objective.

### Image processing, quantification and analysis

To generate distribution of Foxp2 and Pou6f2 cell bodies, 27 representative images across 3 brains were visualized in Imaris. Cell body coordinates were extracted within the SNr using the Spots function with respect to the cerebral aqueduct and normalized to brain section dimensions. Subsequent analyses were performed in MATLAB. Quantification of the colocalization of retrogradely labeled SNr neurons with Foxp2 and Pou6f2 expression was performed in Imaris by segmenting the SNr using the Surface function, identifying fluorescent expression using the Spots function and determining overlap using the Colocalization function. Quantification of RNAscope experiments was performed manually in ImageJ/Fiji. *Pvalb* relative intensity was and normalized to background intensity for each brain section.

Automated reconstructions of slide-scanned images were performed in ImageJ/Fiji^29^ using BrainJ as previously described.^30,31^ Briefly, brain sections were aligned and registered using two-dimensional (2D) rigid-body registration.^32^ A seven-pixel rolling-ball filter was used on all images to reduce background signal and a machine-learning pixel classification approach using Ilastik was used to identify cell bodies and/or neuronal processes.^33^ Several background-subtracted images per fluorescence channel were imported into Ilastik and representative cell bodies, neurites and background fluorescence features were selected to train the algorithm across all images. Probabilistic assignment of image features was continuously checked with a live preview feature to ensure accuracy. The algorithms were then used to generate probability images for each fluorescence channel of each brain section image, and the resulting images were processed for segmentation of cell bodies and neurites. To map the location of these structures to an annotated brain atlas, 3D image registration was performed using Elastix relative to a reference brain.^34^ The coordinates of detected cells and processes were then projected into the Allen Brain Atlas Common Coordinate Framework.^35^ For detection of cells in subdomains of the striatum, detected cells were projected into the Enhanced and Unified Atlas.^36^ Visualizations of the data were performed in ImageJ and Imaris, and subsequent analyses were performed in MATLAB using custom software.

### Nuclear isolation and 10x sequencing of SNr neurons

For 10x sequencing of unlabeled SNr neurons, 9 12-week old C57BL/6JN mice were used. Animals were euthanized in a CO2 chamber and decapitated, after which the brains were dissected out from the skull and submerged in ice-cold aCSF (92 mM NMDG, 2.5 mM KCl, 1.25 mM NaH2PO4, 30 mM NaHCO3, 20 mM HEPES, 25 mM glucose, 5 Na-ascorbate, 3 mM Na-pyruvate, 0.5 mM CaCl2·2H2O, and 10 mM MgSO4·7H2O). Coronal brain slices (400 µm thick) were cut using a vibrating-blade microtome (V1200S, Leica, USA) at 4°C. The SNr subregion was microdissected out in aCSF under a fluorescent dissection microscope (Leica M165FC) and transferred to microcentrifuge tubes. Excess aCSF was aspirated before flash-freezing the tubes on dry ice.

Single nuclei preparations were performed pooling tissue from 3-4 mice, following the Allen Institute’s published V1 protocol.^37^ Briefly, each pooled sample was homogenized in a chilled 2 mL dounce (Sigma D8938) using 3.5 ml of freshly prepared homogenization buffer containing 3.375 ml of NIM (10 mM Tris-HCl, 250 mM sucrose, 25 mM KCl, 5mM MgCl2, 10mM Tris-HCl), 3.5 µl of 100 mM DTT stock (Promega P1171), 70 µl of 50X Protease inhibitor cocktail (Promega G6521), 17.5 µl of 40 U/µl RNAsin Plus (Promega N2615), and 35 µl of 10% Triton X-100 (Sigma T8787). The homogenate was strained through a 30 µm cell strainer (Miltenyi Biotec 130-098-458) and centrifuged at 900 x g for 10 minutes at 4°C. After centrifugation, the pellet was resuspended in 500 µl of freshly prepared blocking buffer containing 410 µl nuclease-free water, 50 µl 10X PBS, 40 µl of 10% OmniPur BSA (Sigma 2905), and 2.5 µl of 40 U/µl RNAsin Plus. The nuclear resuspension as subsequently incubated on ice for 15 minutes. The suspension was labeled with Mouse anti-NeuN-488 to enrich for neuronal cells (1:500, Millipore Sigma), incubated on ice for 30 minutes, then centrifuged at 400 x g for 5 minutes. The supernatant was resuspended in 500 µl of blocking buffer, then labeled with 0.5 µl of DAPI. Prior to DAPI labeling, 5 µl of nuclear suspension was removed and labeled with 5 µl of Trypan Blue and loaded into a disposable hemocytometer slide (Invitrogen C10228) for quantification of yield and visual inspection of nuclear quality.

Fluorescence-activated nuclei sorting (FANS) of single nuclei was performed using the Beckman Coulter MoFlo Astrios EQ. Cell analysis and sorting were performed on a Beckman Coulter MoFlo Astrios EQ. These experiments required the use of the 405 nm (80 mW) laser for fluorescent excitation of the DAPI labeled nuclei and the 488 nm (200 mW) laser for forward, side scatter detection and GFP excitation. The sample and collection area were cooled to ∼ 7° C. DAPI was detected with a 448 /59 band-pass filter. The PMT was set at 341V with an amplifier gain of 3. GFP was detected with a 513 /26 band-pass filter. The PMT was set at 360V with an amplifier gain of 3. The PMT was set at 310V with an amplifier gain of 4. DAPI^+^/GFP^+^ nuclei were collected into an Eppendorf tube containing blocking buffer.

### 10x library prep and sequencing

Single-nucleus cDNA libraries were constructed according to the user guide of Chromium Single Cell 3’ Reagents Kit v3 (10× Genomics, Pleasanton, CA). Libraries were sequenced on the Illumina NovaSeq 6000 and sequencing reads were processed and aligned to the mm10 transcriptome using Cell Ranger 5.0.1. 3999 cells were captured with 99874 mean reads per nucleus and 4304 median genes per nucleus.

### Analysis of 10x RNAseq dataset

The Seurat (v4.3) package was used to perform clustering analysis of the 10x snRNASeq dataset as described.^38^ 1403 low quality samples (nFeature < 2500) were excluded, yielding 2596 samples for subsequent analysis. The dataset was normalized and 4000 variable genes were identified before scaling and performing PCA. The optimal dimensionality for clustering was determined using the ‘Elbowplot’ function. The optimal resolution for clustering was determined by performing iterative clustering with 9 levels of resolution (0.2-1.0, 0.1 increments) using the ‘Clustree’ function.^39^ Clustering was performed using 20 PCs and a resolution of 0.4, yielding 12 clusters. 1 cluster (105 nuclei) corresponding to glia was identified based on expression of known glial markers (Apo3, Olig2, Sox10, Mag, Mbp). 2 clusters (501 samples total) corresponding to dopamine neurons were identified based on expression of Th and Slc6a3. 2 clusters (491 samples total) corresponding to glutamatergic neurons were identified based on expression of Slc17a6 and Slc17a7. The remaining 7 clusters (1499 samples total) were found to be GABAergic based on expression of Gad1 and Gad2 and presumed to correspond to SNr neurons. Cluster specific markers were identified using the ‘FindMarkers’ function.

This processed dataset was subsequently mapped to the Allen Brain Cell Atlas taxonomy using MapMyCells.^12^ 1432 of 1499 samples mapped to ABCA taxonomies with 10 or more cells per taxonomy label. 31 cells were removed from analysis that mapped to ABCA subclass and supertype taxonomies corresponding to less than 10 cells and 36 cells were removed that mapped to ABCA cluster taxonomies corresponding to less than 10 cells. All RNAseq plots were generated using R statistical computing environment (R Core Team (2017). R: A language and environment for statistical computing. R Foundation for Statistical Computing, Vienna, Austria. URL https://www.R-project.org/)

### Nuclear isolation and plate-based sequencing after retrograde labeling

For plate-based sequencing of SNr neurons retrogradely labeled by hindbrain injection of AAVretro-hSyn-H2B-mCherry, 24 12-week old C57BL/6JN mice were used (3 mice for LSC, 9 for MSC, 3 for IC, 3 for DR, 3 for PNo, 3 for Med). Tissue was prepared and nuclei were isolated as before without NeuN antibody labeling for neuronal enrichment. FANS sorting was performed as before using a 405 nm (80 mW) laser for fluorescent excitation of the DAPI labeled nuclei and the 561 nm (200 mW) laser for excitation of mCherry. mCherry was detected with a 614 /20 band-pass filter. DAPI^+^/mCherry^+^ nuclei were collected and sorted into 96-well plates preloaded with a lysis solution (6 µl per well) including 0.2% Triton X-100, 1 U/ul SuperaseIN (Invitrogen), 1 mM dNTPs and RT primer. Individual nuclei were sorted into the 96-well plates on “Single Cell” mode with a drop envelope of “1”. Sheath pressure was set at 28 PSI. Sample pressure was kept below 28.5 PSI. A 100 µm nozzle was used. Cells were run at an event rate between 2000 – 5000 cells / second. For cell sorting experiments a droplet drive frequency of 49.6 KHz was used.

For LSC, 4 plates containing 384 neurons were collected; for MSC, 2 plates containing 192 neurons; for IC, 3 plates containing 288 neurons; for DR, 5 plates containing 480 neurons; for PNo, 4 plates containing 384 neurons; for Med, 4 plates containing 384 neurons. After sorting, 96-well plates were briefly centrifuged, flash frozen on dry ice and stored at -80°C prior to library preparation.

### Plate-based library prep and sequencing

Primer annealing was performed by incubating plates on a thermocycler at 72°C for 3 minutes, followed by hold at 4°C. A reverse transcription reaction mix (7.5 µl) was added to each well. Each 7.5 µl of reaction mix contained 40U Maxima H Minus Reverse Transcriptase (ThermoFisher), 4U SupernaseIN, 7.5% PEG and 1uM TSO and 1X Maxima Buffer. Reverse transcription was performed by incubating plates on a thermocycler at 42°C for 90 minutes, followed by 10 cycles of 50°C for 2 minutes and 42°C for 2 minutes. Reverse transcription was stopped by heating at 75°C for 10 minutes, followed by hold at 4°C. An ExoI digestion mix (0.22 U/ µl) was subsequently added to each well to remove excess primer or singled stranded products. ExoI digestion was performed by incubating plates on a thermocycler at 37°C for 30 minutes, 85°C for 15 minutes and 75°C for 30 seconds, followed by hold at 4°C. Pooling was performed on the Biomek 4000 (Beckman Coulter, Brea California). cDNA cleanup was performed using Silane beads (ThermoFisher) suspended in RLT Plus buffer (Qiagen) and ethanol. Bead incubation was performed at room temperature for 20 minutes followed by placement on a magnetic stand. The supernatant was removed and the beads were washed twice with 80% ethanol before being returned to the magnet and allowed to air dry for 10 minutes. The beads were resuspended in 50 µl of nuclease-free water, incubated for 2 minutes and returned to the magnet. The supernatant was then removed, transferred to a clean Eppendorf tube and then split into two tubes for cDNA amplification.

cDNA amplification was performed using a master mix containing 25 µl of 2X Kapa HotStart Mix (Roche) per reaction and 2 µl of 5 µM SMART PCR primer (Takara Bio). Amplification was performed by incubating tubes on a thermocycler at 98°C for 3 minutes followed by 18 cycles of 98°C for 20 seconds, 67°C for 15 seconds and 72°C for 5 minutes. The reaction was then maintained at 72°C for 5 minutes, followed by hold at 4°C. The two reactions were pooled in one Eppendorf tube and 70 µl of AMPureXP beads (Beckman Coulter) were added to the sample and incubated for 5 minutes at room temperature. The sample was placed on a magnet and the supernatant was removed. The beads were then washed twice with 80% ethanol, returned to the magnet and dried for 5 minutes. The beads were then resuspended in 25 µl of nuclease-free water, incubated for 2 minutes and placed on the magnet for removal of the supernatant to a new tube. The quality and quantity of cDNA was examined using an Agilent Bioanalyzer and Qubit Fluorometer (ThermoFisher).

Sequencing libraries were prepared using Nextera XT Library Prep Kit (Illumina). 0.6 ng of cDNA was used as input and tagmentation and neutralization was performed as directed. Indexing PCR was performed as directed using 10 cycles by adding 5 µl of 2 µM custom i5 primer, 5 µl N7 Nextera primer and 15 µl of NPM. PCR was followed by 2 AMPureXP cleanups as above, first with 0.6 beads:sample ratio and then with 1:1 beads:sample. The final volume was eluted in 20 µl of nuclease-free water. QC was repeated as before. Libraries were sequenced using an Illumina NextSeq 500/550. Sequences were de-multiplexed to generate a fastq file. Reads were aligned to annotated mRNAs in the mouse genome (UCSC, Mus musculus assembly mm10) and the read count for each gene was calculated using STAR aligner.^40^

### Analysis of integrated RNAseq datasets

The Seurat (v4.3) package was used to perform integration and clustering analysis of the combined 10x and projection-tagged datasets. 100 low quality samples (nFeature < 2500) were excluded from the projection-tagged dataset, yielding 2012 samples for subsequent analysis. After merging the projection-tagged dataset with the 2596 samples from the 10x dataset, each dataset was normalized and 4000 variable genes were identified in each. The datasets were integrated (anchor.features = 3000, dims = 1:30), yielding 27748 features across a total of 4608 samples. After scaling and performing PCA, the optimal dimensionality and resolution for clustering was performed as before. Clustering was performed using 20 PCs and a resolution of 0.3, yielding 17 clusters. 1 cluster (140 samples) corresponding to glia was identified. 1 cluster (501 samples) corresponding to dopamine neurons was identified, nearly all deriving from the 10x dataset. 6 clusters (918 samples) corresponding to glutamate neurons were identified, deriving equally from the 10x and projection-tagged datasets. The remaining 9 clusters (3049 samples total, 1507 from the 10x dataset and 1542 from the projection-tagged dataset) were GABAergic and presumed to correspond to SNr neurons. Of the 1542 GABAergic SNr neurons from the projection-tagged dataset, 280 were from LSC, 115 from MSC, 171 from IC, 416 from DR, 302 from PNo, and 258 from Med. The processed dataset was subsequently mapped to the ABCA taxonomy using MapMyCells. 2965 of 3049 samples mapped to ABCA taxonomies with 10 or more cells per taxonomy label. 84 samples were removed from analysis that mapped to ABCA subclass, supertype or cluster taxonomies corresponding to less than 10 cells.

### Analysis of human RNAseq dataset

The Harmony (v1.2) package^41^ was used to perform integration and the Seurat (v4.3) package was used to perform clustering analysis of 91479 human non-dopaminergic neurons from control, Parkinson’s Disease and Lewy Body Dementia donors sequenced in a previously published dataset.^15^ 12429 low quality samples (nFeature < 2500) were excluded, yielding 38594 features across 79050 samples. For analysis of human SNr neurons from control donors, 39110 samples corresponding to 8 control donors were subsetted and normalized. Variable features were identified before scaling and performing PCA. Integration was performed across donor ID using Harmony. The optimal resolution for clustering was performed as before and clustering was performed using 20 PCs and a resolution of 0.4, yielding 23 clusters. 4 clusters (12640 samples) of low-quality samples were excluded based on lack of expression of known neurotransmitter genes, expression of immediate release genes (FOS, EGR1, NR4A1, DUSP1, JUN) or nCount < 10000. 11 clusters (17348 samples) corresponding to glutamatergic neurons were identified based on expression of SLC17A6 and SLC17A7. The remaining 8 clusters (9122 samples) were found to be GABAergic based on expression of GAD1 and GAD2.

For analysis of human SNr neurons from control and Parkinson’s Disease donors, 39110 samples from 8 control donors and 34779 samples from 7 PD donors were merged and normalized. Variable features were identified before scaling and performing PCA. Integration was performed across disease state using Harmony. Clustering was performed using 20 PCs and a resolution of 0.2, yielding 22 clusters. 2 low quality clusters and 14 glutamatergic clusters were excluded. The remaining 6 clusters (13910 samples) were GABAergic (9307 control samples, 4603 PD samples) and used for subsequent differential gene expression analysis.

### Statistical analysis

All sample sizes, statistical tests and significance values are reported in the Results or figure legends. Data is mean ± standard error of mean (SEM). Statistical tests were performed using GraphPad and significance is defined as p < 0.05.

## FIGURE LEGENDS

**Figure S1:**
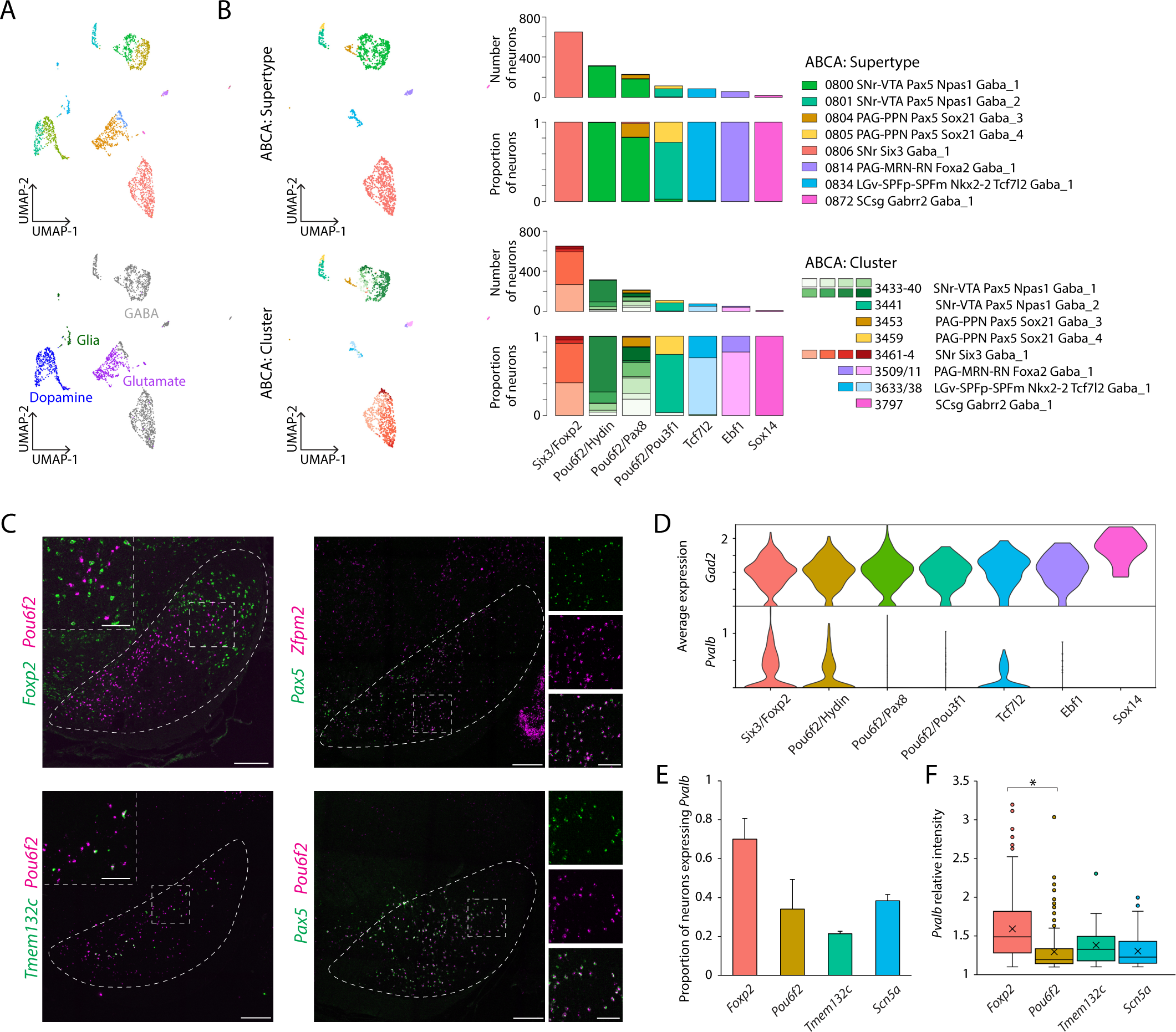
Characterization of SNr cell types, related to Figure 1. **(A)** UMAP representation of 2596 nuclei isolated after SNr microdissection, colored by assigned cluster (top) and cell type label (bottom). **(B)** SNr cluster identity mapped to Allen Brain Cell Atlas mouse taxonomy with MapMyCells. UMAP representations of SNr GABAergic neurons from 1B colored by ABCA supertype (top left) or ABCA cluster (bottom left). Number and proportion of neurons per cluster mapping to corresponding ABCA label (right). **(C)** RNAscope reveals marker distribution within SNr. Scale bars are 250µm, inset and rightside panel scalebars are 100µm. **(D)** Violin plots showing average expression of *Gad2* and *Pvalb* per SNr cluster. **(E)** Proportion of *Foxp2^+^*, *Pou6f2^+^*, *Tmem132c^+^* and *Scn5a^+^* neurons labeled by RNAscope that colocalize with *Pvalb*. N = 3 mice, 6 slides, *Foxp2*: 279 neurons, *Pou6f2*: 335 neurons, *Tmem132c*: 150 neurons, *Scn5a*: 187 neurons. **(F)** Relative intensity of *Pvalb* expression in *Foxp2^+^*, *Pou6f2^+^*, *Tmem132c^+^* and *Scn5a^+^* neurons colocalizing with *Pvalb*. N = 3 mice, 6 slides, *Foxp2*: 225 neurons, *Pou6f2*: 153 neurons, *Tmem132c*: 31 neurons, *Scn5a*: 71 neurons. Data are represented as mean ± SEM. *: p < 0.0001.

**Figure S2.**
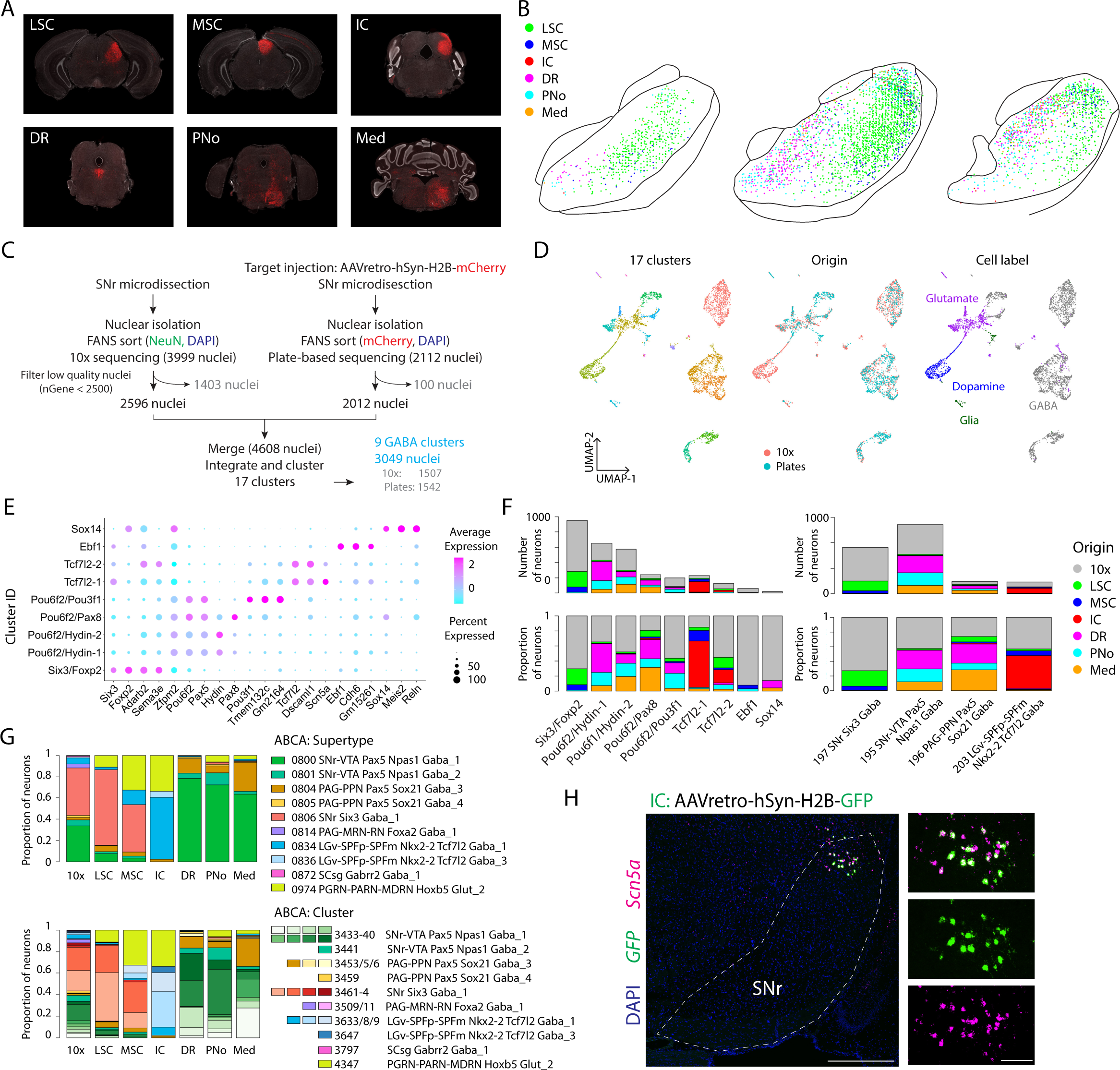
Characterization of projection-defined SNr cell types, related to Figure 2. **(A)** Six SNr target sites injected with AAVretro-hSyn-H2B-mCherry. **(B)** Reconstruction of coordinate locations of SNr neurons retrogradely labeled by injection of AAVretro-hSyn-H2B-mCherry in each of six target sites. **(C)** Experimental schematic for integrated analysis of 10x SNr profiling and single-nucleus RNAseq profiling of retrogradely labeled SNr neurons. **(D)** UMAP representation of 4608 SNr neurons from integrated cluster analysis of whole SNr 10x (2596 neurons) and projection-tagged plate-based RNAseq (2012 neurons), colored by assigned cluster, origin or cell type label. **(E)** Dotplot showing expression of selected marker genes across clusters. **(F)** Stacked barplot showing the number and proportion of SNr neurons per cluster (left) and mapped ABCA subclass (right) originating from the 10x dataset or each projection target. **(G)** Proportion of 10x or retrogradely labeled SNr neurons corresponding to a given ABCA supertype (top) or ABCA cluster (bottom). **(H)** SNr neurons retrogradely labeled by IC injection of AAVretro-hSyn-H2B-GFP colocalize with *Scn5a*. Scalebars are 500µm (left) and 100µm (right panels).

**Figure S3.**
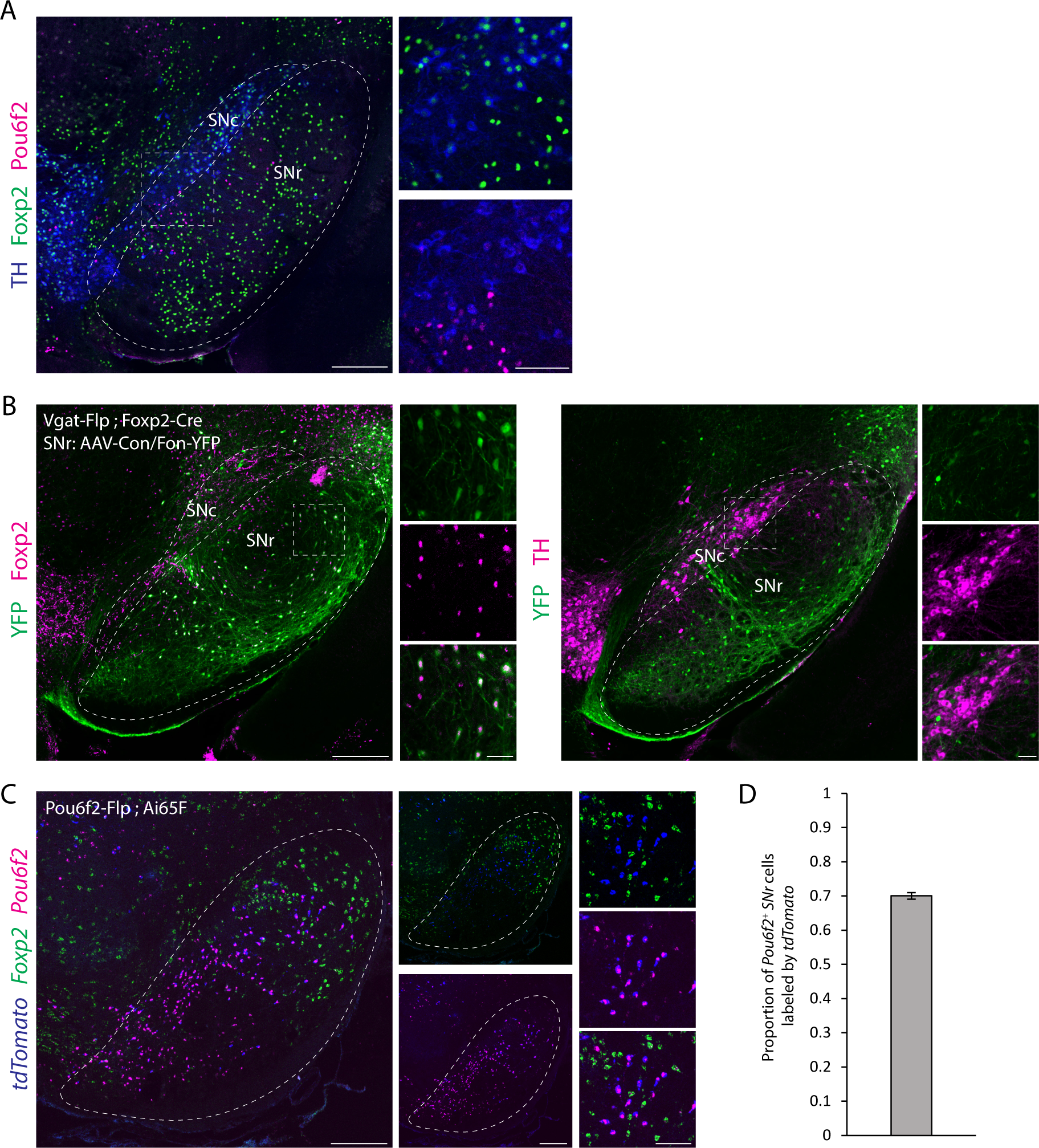
Validation of Foxp2 and Pou6f2 subclass labeling strategies, related to Figure 3. **(A)** TH^+^ dopamine neurons in the substantia nigra pars compacta (SNc) express Foxp2. SNc adjacent Pou6f2^+^ SNr neurons do not express TH. Scalebar is 250µm (left) and 100µm (right) **(B)** Injection of AAV-Con/Fon-YFP into the SNr of Vgat-Flp;Foxp2-Cre mice labels TH^-^/Foxp2^+^ cells in SNr but not TH^+^ cells in SNc. Scalebar is 250µm (large panels) and 50µm (small panels). **(C)** Validation of Pou6f2-Flp mouse line with RNAscope. *TdTomato^+^* cells in Pou6f2-Flp;Ai65F mice colocalize with endogenous expression of *Pou6f2* but not *Foxp2*. Scalebars are 250µm (left and middle panels) and 100µm (right panels). **(D)** Proportion of SNr neurons expressing endogenous *Pou6f2* by RNAscope that are labeled by *tdTomato* in Pou6f2-Flp;Ai65 mice. N = 3 mice, 6 sections, 3870 *Pou6f2^+^* neurons. Data are represented as mean ± SEM.

**Figure S4.**
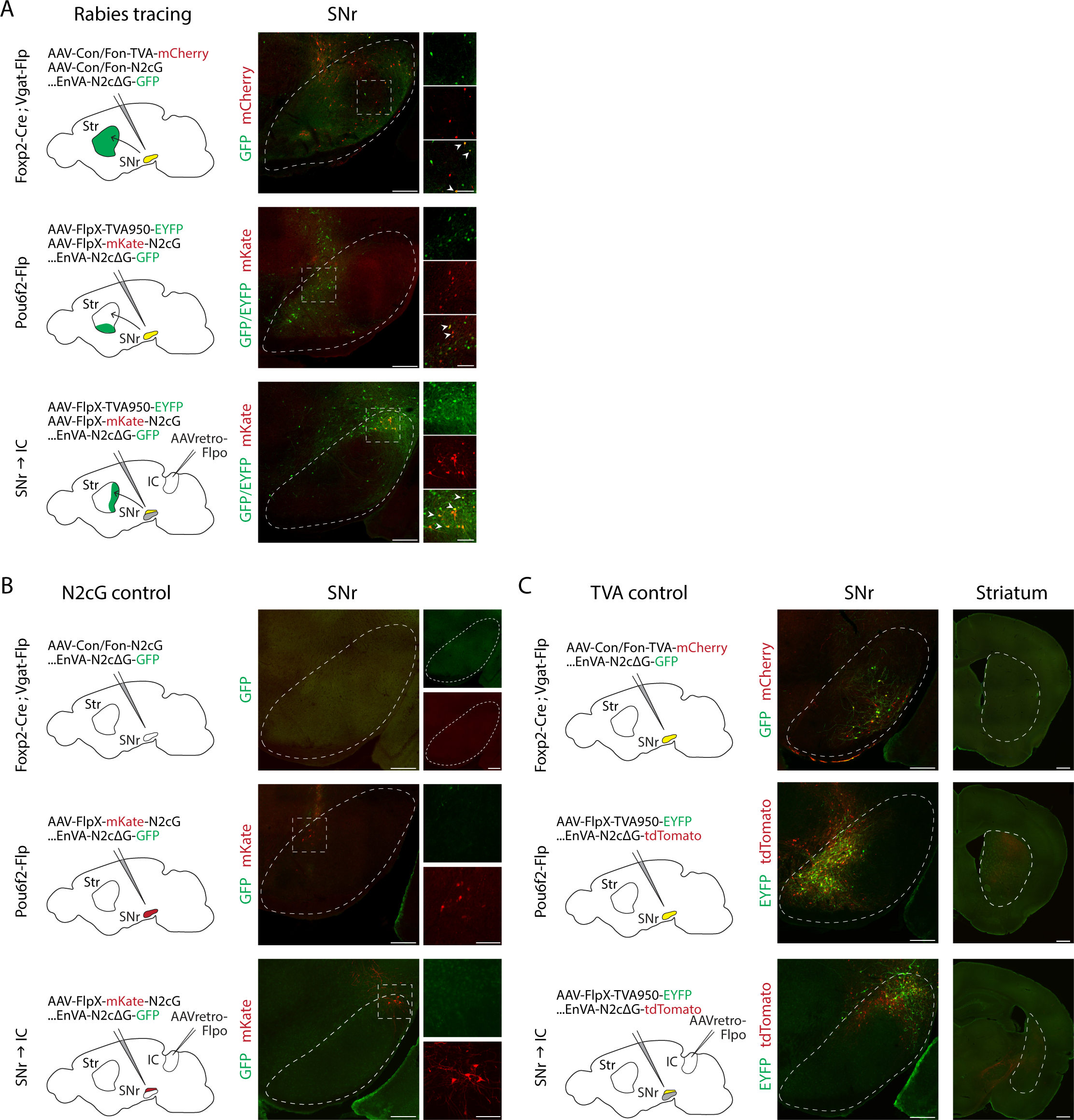
Validation of rabies labeling strategy, related to Figure 4. **(A)** Starter cells in SNr labeled with rabies virus after injection of TVA and N2cG helper viruses. Scalebars are 250µm (left) and 100µm (right). **(B)** Rabies virus does not label cells in SNr after injection of N2cG helper virus without TVA helper virus. Scalebars are 250µm (left panels and top right small panels) and 100µm (middle and bottom small panels). **(C)** Rabies virus labels cells in SNr after injection of TVA helper virus but fails to label presynaptic input cells in striatum. Scalebars are 250µm (left) and 500µm (right).

**Figure S5.**
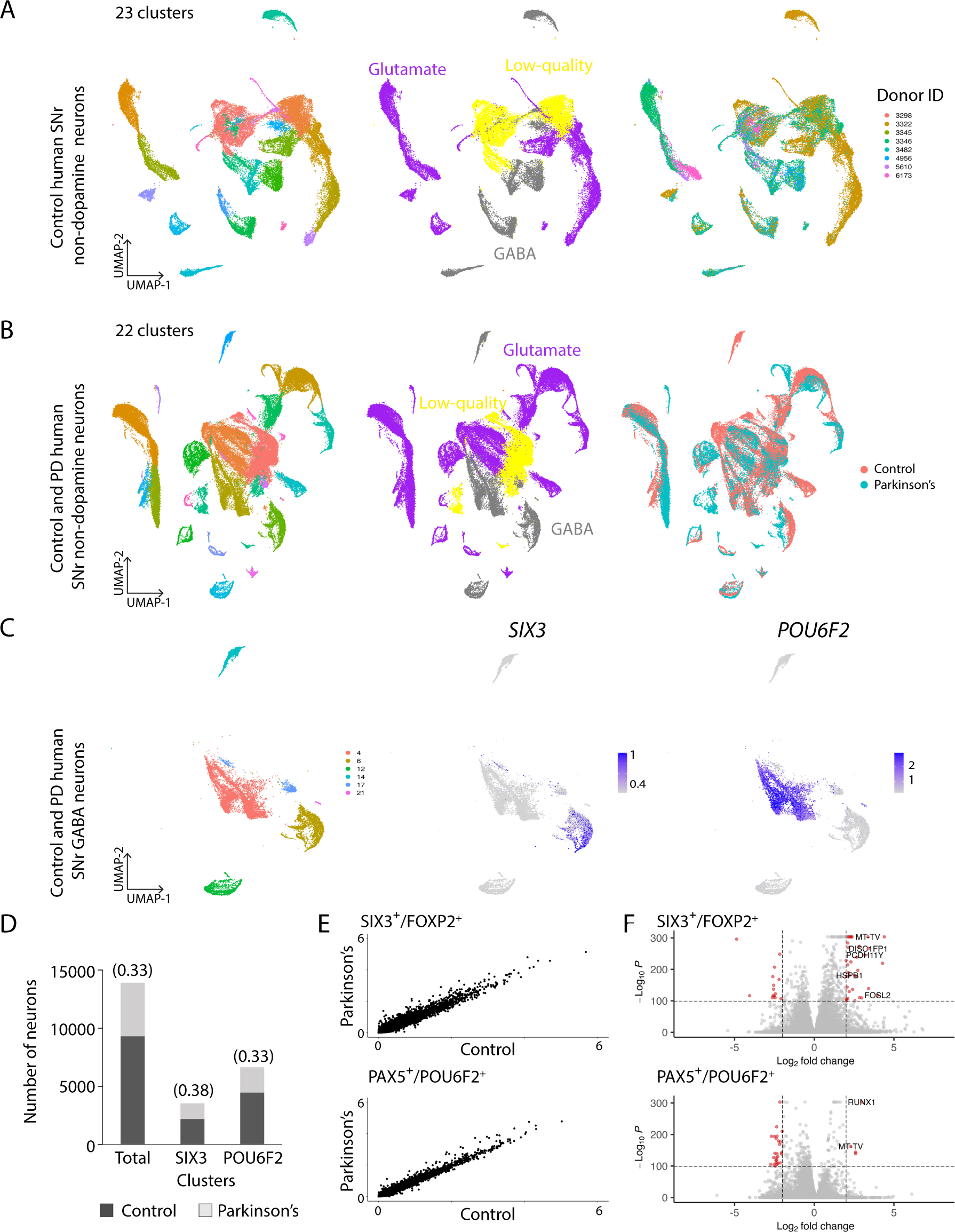
Single-cell transcriptomic profiling of control and Parkinson’s human SNr neurons, related to Figure 5. **(A)** UMAP representation of 39110 human non-dopaminergic midbrain neurons from 8 control donors, colored by cluster (left), cell type label (middle) and donor ID (right). **(B)** UMAP representation of 73889 human non-dopaminergic midbrain neurons from 8 control donors (39110 neurons) and 7 Parkinson’s Disease donors (34779 neurons), colored by cluster (left), cell type label (middle) and disease state (left). **(C)** UMAP representation of 13910 human GABAergic SNr neurons (9307 control neurons and 4603 PD neurons) subsetted from (B) and feature plots showing relative expression of *SIX3* and *POU6F2*. **(D)** Number of neurons in the SIX3^+^/FOXP2^+^ and PAX5^+^/POU6F2^+^ clusters deriving from control or PD donors. Proportion from PD donors shown in parentheses. **(E)** Average gene expression of control and PD neurons within SIX3^+^/FOXP2^+^ clusters (top) and PAX5^+^/POU6F2^+^ clusters (bottom). **(F)** Volcano plots of genes up- or downregulated in PD compared to control within SIX3^+^/FOXP2^+^ clusters (top) and PAX5^+^/POU6F2^+^ clusters (bottom). Genes shown in red have fold change > 2 and p < 10e-100.

**Table S1.** Select SNr target brains regions and their acronyms, related to Figure 3.

|  |  |
| --- | --- |
| PPN | Pedunculo pontine nucleus |
| ZI | Zona incerta |
| PAG | Periaqueductal gray |
| DR | Dorsal nucleus raphe |
| CUN | Cuneiform nucleus |
| RN | Red nucleus |
| SCig | Superior colliculus, motor related, intermediate gray layer |
| SCiw | Superior colliculus, motor related, intermediate white layer |
| SCdg | Superior colliculus, motor related, deep gray layer |
| ICe | Inferior colliculus, external nucleus |
| ICc | Inferior colliculus, central nucleus |
| ICd | Inferior colliculus, dorsal nucleus |
| PRNr | Pontine reticular nucleus |
| PRNc | Pontine reticular nucleus, caudal part |
| GRN | Gigantocellular reticular nucleus |
| PARN | Parvocellular reticular nucleus |
| IRN | Intermediate reticular nucleus |
| VAL | Ventral anteriorlateral complex of the thalamus |
| VM | Ventral medial nucleus of the thalamus |
| MD | Mediodorsal nucleus of thalamus |
| CM | Central medial nucleus of the thalamus |
| PF | Parafascicular nucleus |
| RT | Reticular nucleus of the thalamus |
| PIL | Posterior intralaminar thalamic nucleus |
| PoT | Posterior triangular thalamic nucleus |
| MGv | Medial geniculate complex, ventral part |
| MGm | Medial geniculate complex, medial part |
| MGd | Medial geniculate complex, dorsal part |

**Table S2.**
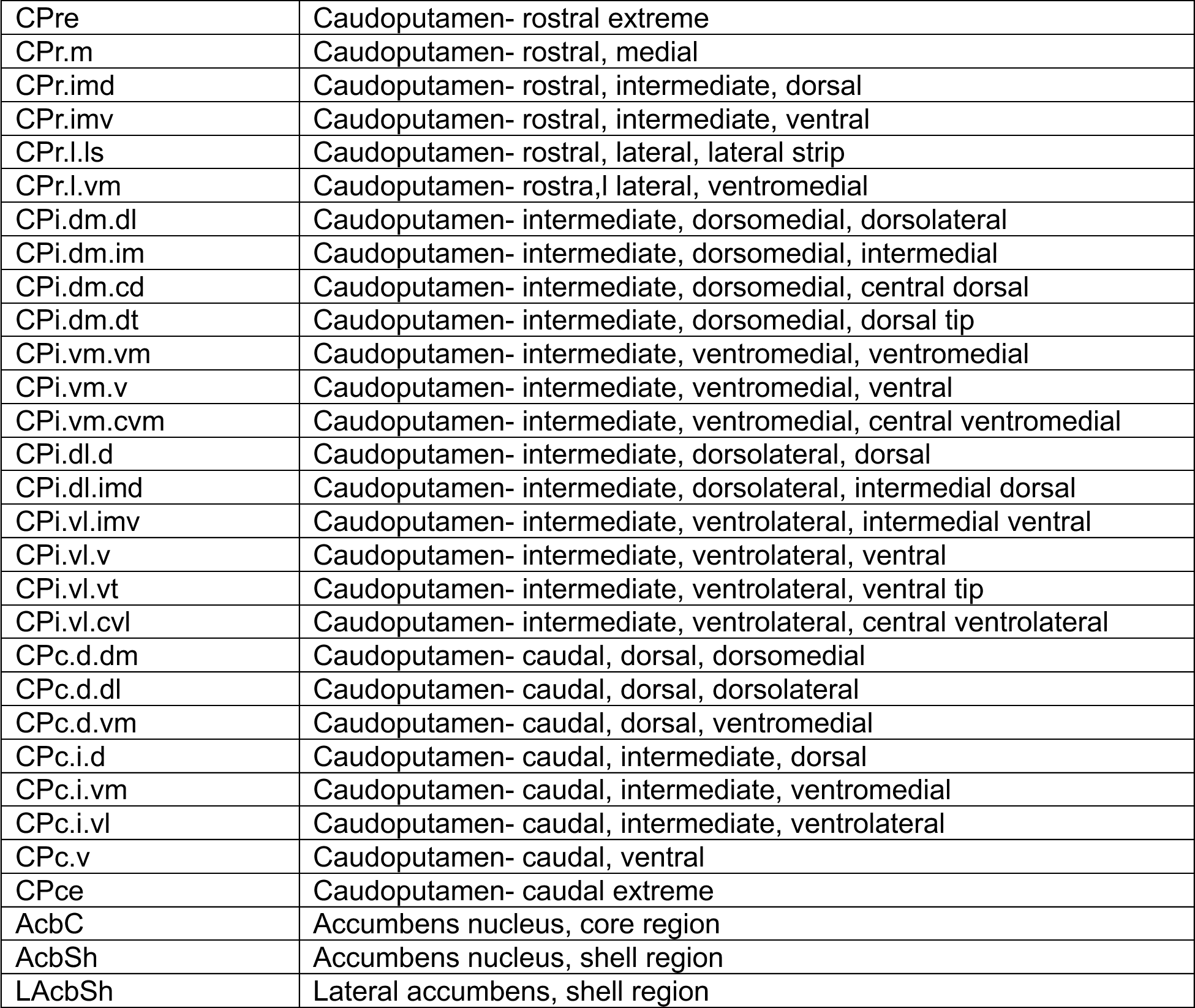
Striatum subdomains and their acronyms, related to Figure 4.

|  |  |
| --- | --- |
| CPre | Caudoputamen- rostral extreme |
| CPr.m | Caudoputamen- rostral, medial |
| CPr.imd | Caudoputamen- rostral, intermediate, dorsal |
| CPr.imv | Caudoputamen- rostral, intermediate, ventral |
| CPr.l.ls | Caudoputamen- rostral, lateral, lateral strip |
| CPr.l.vm | Caudoputamen- rostra,l lateral, ventromedial |
| CPi.dm.dl | Caudoputamen- intermediate, dorsomedial, dorsolateral |
| CPi.dm.im | Caudoputamen- intermediate, dorsomedial, intermedial |
| CPi.dm.cd | Caudoputamen- intermediate, dorsomedial, central dorsal |
| CPi.dm.dt | Caudoputamen- intermediate, dorsomedial, dorsal tip |
| CPi.vm.vm | Caudoputamen- intermediate, ventromedial, ventromedial |
| CPi.vm.v | Caudoputamen- intermediate, ventromedial, ventral |
| CPi.vm.cvm | Caudoputamen- intermediate, ventromedial, central ventromedial |
| CPi.dl.d | Caudoputamen- intermediate, dorsolateral, dorsal |
| CPi.dl.imd | Caudoputamen- intermediate, dorsolateral, intermedial dorsal |
| CPi.vl.imv | Caudoputamen- intermediate, ventrolateral, intermedial ventral |
| CPi.vl.v | Caudoputamen- intermediate, ventrolateral, ventral |
| CPi.vl.vt | Caudoputamen- intermediate, ventrolateral, ventral tip |
| CPi.vl.cvl | Caudoputamen- intermediate, ventrolateral, central ventrolateral |
| CPc.d.dm | Caudoputamen- caudal, dorsal, dorsomedial |
| CPc.d.dl | Caudoputamen- caudal, dorsal, dorsolateral |
| CPc.d.vm | Caudoputamen- caudal, dorsal, ventromedial |
| CPc.i.d | Caudoputamen- caudal, intermediate, dorsal |
| CPc.i.vm | Caudoputamen- caudal, intermediate, ventromedial |
| CPc.i.vl | Caudoputamen- caudal, intermediate, ventrolateral |
| CPc.v | Caudoputamen- caudal, ventral |
| CPce | Caudoputamen- caudal extreme |
| AcbC | Accumbens nucleus, core region |
| AcbSh | Accumbens nucleus, shell region |
| LAcbSh | Lateral accumbens, shell region |

## Key Resource Table

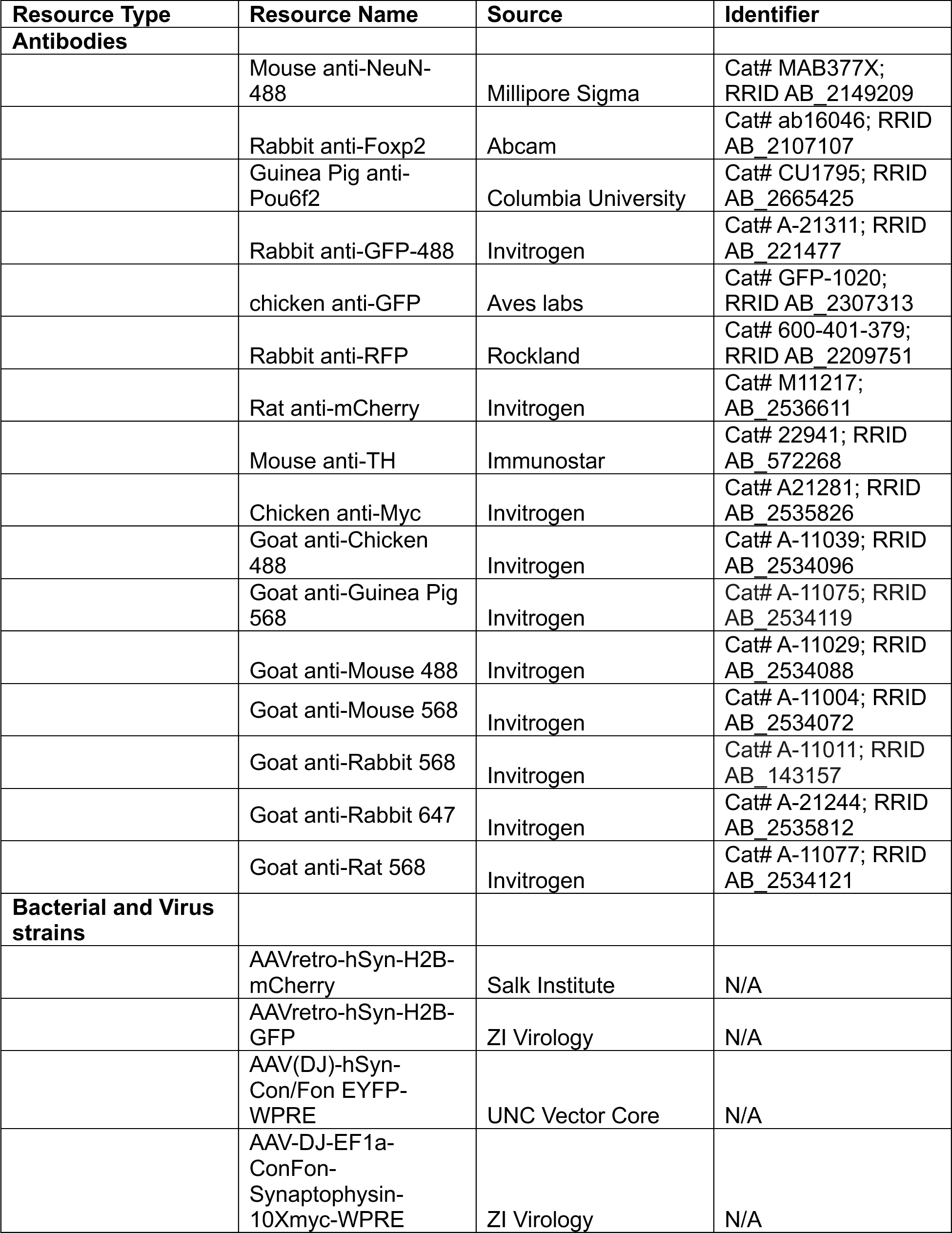
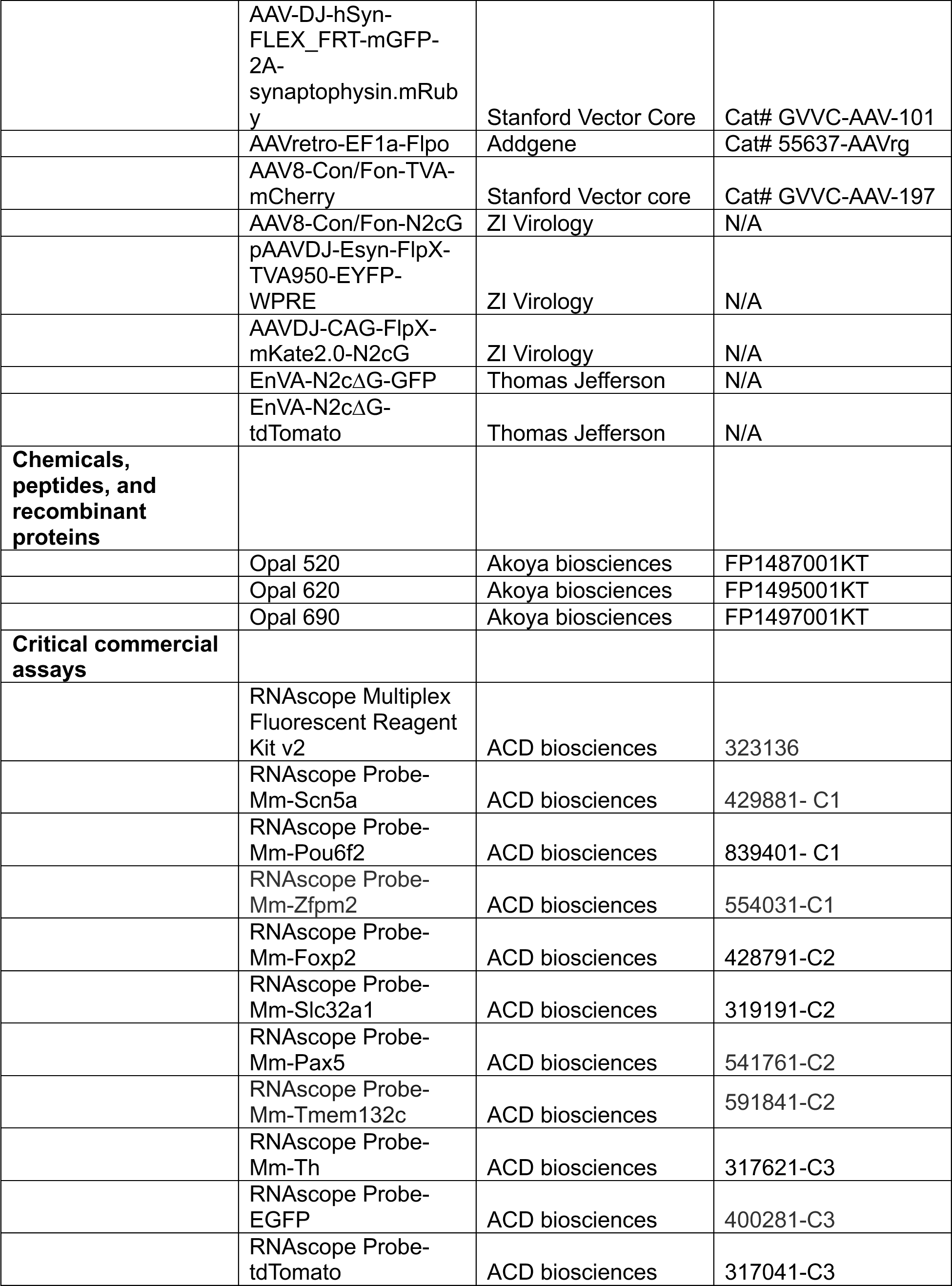
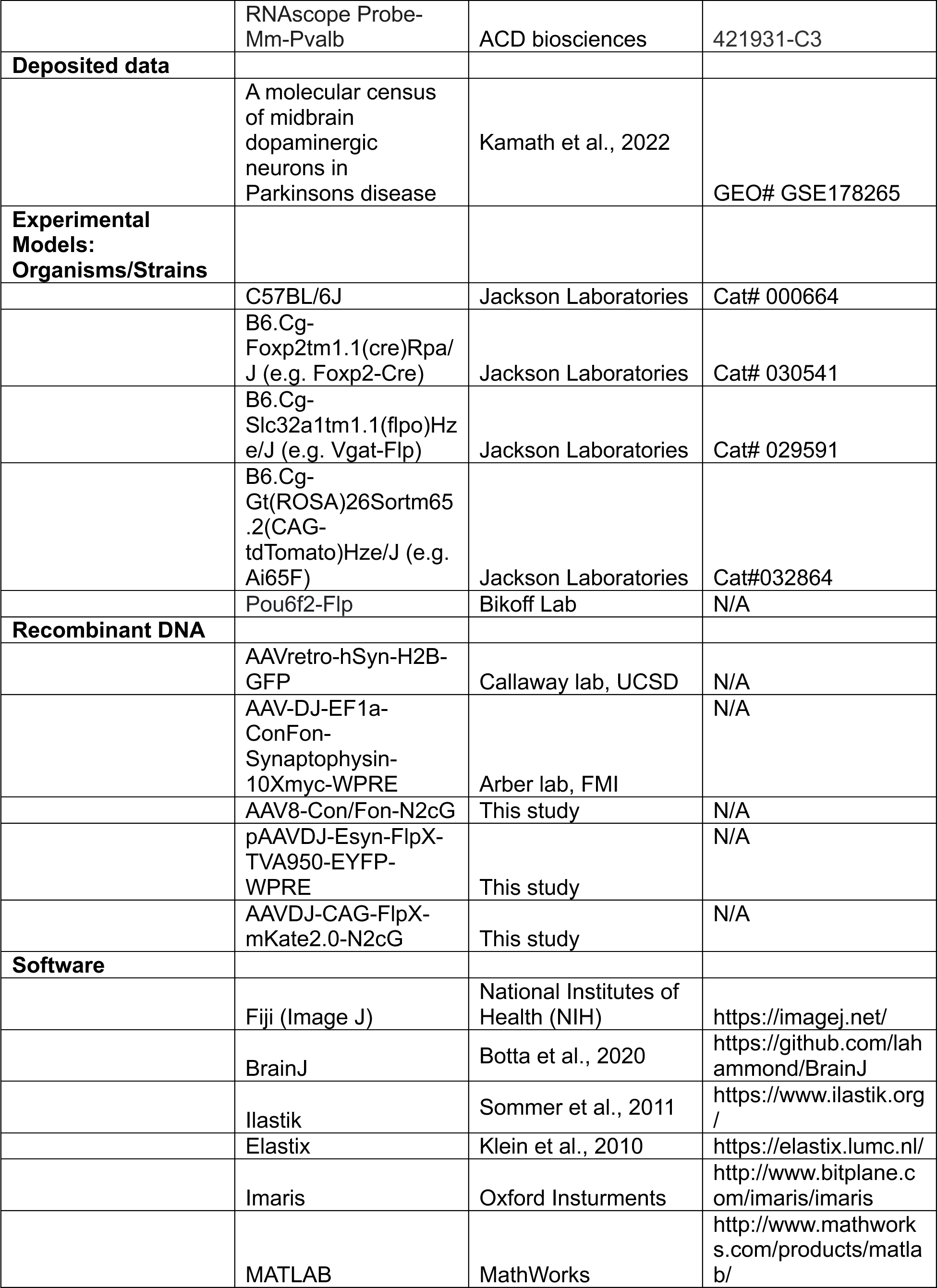
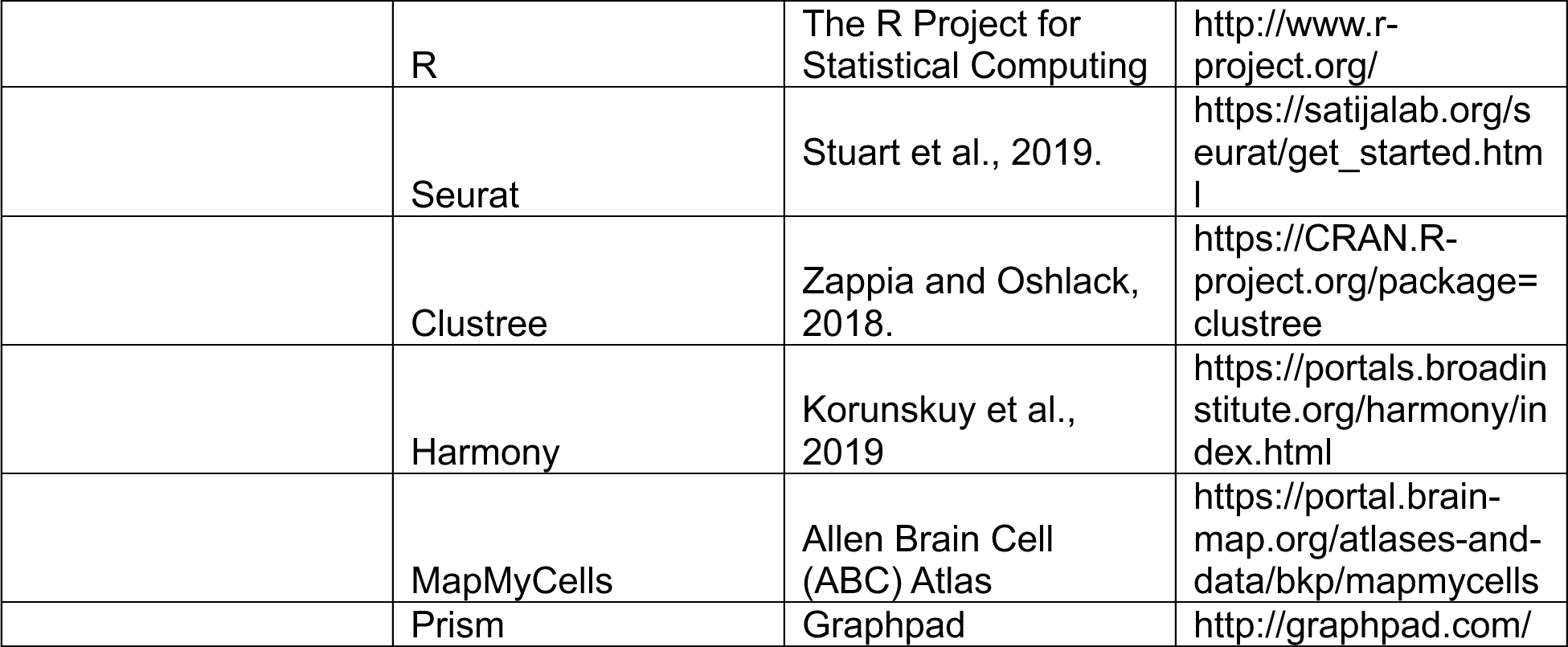

## REFERENCES

1. Arber, S., and Costa, R.M. (2022). Networking brainstem and basal ganglia circuits for movement. Nature Reviews Neuroscience 23, 342–360. 10.1038/s41583-022-00581-w.

2. Hintiryan, H., Foster, N.N., Bowman, I., Bay, M., Song, M.Y., Gou, L., Yamashita, S., Bienkowski, M.S., Zingg, B., Zhu, M., et al. (2016). The mouse cortico-striatal projectome. Nature Neuroscience 19, 1100–1114. 10.1038/nn.4332.

3. Foster, N.N., Barry, J., Korobkova, L., Garcia, L., Gao, L., Becerra, M., Sherafat, Y., Peng, B., Li, X., Choi, J., et al. (2021). The mouse cortico – basal ganglia – thalamic network. Nature 598. 10.1038/s41586-021-03993-3.

4. Lee, J., Wang, W., and Sabatini, B.L. (2020). Anatomically segregated basal ganglia pathways allow parallel behavioral modulation. Nature Neuroscience 23, 1388–1398. 10.1038/s41593-020-00712-5.

5. McElvain, L.E., Chen, Y., Moore, J.D., Brigidi, G.S., Bloodgood, B.L., Lim, B.K., Costa, R.M., and Kleinfeld, D. (2021). Specific populations of basal ganglia output neurons target distinct brain stem areas while collateralizing throughout the diencephalon. Neuron, 1–18. 10.1016/j.neuron.2021.03.017.

6. Lahti, L., Haugas, M., Tikker, L., Airavaara, M., Voutilainen, M.H., Anttila, J., Kumar, S., Inkinen, C., Salminen, M., and Partanen, J. (2016). Differentiation and molecular heterogeneity of inhibitory and excitatory neurons associated with midbrain dopaminergic nuclei. Development (Cambridge) 143, 516–529. 10.1242/dev.129957.

7. Madrigal, M.P., Moreno-Bravo, J.A., Martínez-López, J.E., Martínez, S., and Puelles, E. (2016). Mesencephalic origin of the rostral Substantia nigra pars reticulata. Brain Structure and Function 221, 1403–1412. 10.1007/s00429-014-0980-9.

8. Morello, F., Borshagovski, D., Survila, M., Tikker, L., Sadik-Ogli, S., Kirjavainen, A., Estartús, N., Knaapi, L., Lahti, L., Törönen, P., et al. (2020). Molecular Fingerprint and Developmental Regulation of the Tegmental GABAergic and Glutamatergic Neurons Derived from the Anterior Hindbrain. Cell Reports 33. 10.1016/j.celrep.2020.108268.

9. Achim, K., Peltopuro, P., Lahti, L., Li, J., Salminen, M., and Partanen, J. (2012). Distinct developmental origins and regulatory mechanisms for GABAergic neurons associated with dopaminergic nuclei in the ventral mesodiencephalic region. Development (Cambridge) 139, 2360–2370. 10.1242/dev.076380.

10. Saunders, B.T., Richard, J.M., Margolis, E.B., and Janak, P.H. (2018). Dopamine neurons create Pavlovian conditioned stimuli with circuit-defined motivational properties. Nature Neuroscience 21, 1072–1083. 10.1038/s41593-018-0191-4.

11. Allen Brain Cell Atlas (RRID:SCR_024440) https://portal.brain-map.org/atlases-and-data/bkp/abc-atlas.

12. Yao, Z., Van Velthoven, C.T.J., Kunst, M., Zhang, M., McMillen, D., Lee, C., Jung, W., Goldy, J., Abdelhak, A., Aitken, M., et al. (2023). A high-resolution transcriptomic and spatial atlas of cell types in the whole mouse brain. Nature 624, 317–332. 10.1038/s41586-023-06812-z.

13. Zhang, M., Pan, X., Jung, W., Halpern, A.R., Eichhorn, S.W., Lei, Z., Cohen, L., Smith, K.A., Tasic, B., Yao, Z., et al. (2023). Molecularly defined and spatially resolved cell atlas of the whole mouse brain. Nature 624, 343–354. 10.1038/s41586-023-06808-9.

14. La Manno, G., Gyllborg, D., Codeluppi, S., Nishimura, K., Salto, C., Zeisel, A., Borm, L.E., Stott, S.R.W., Toledo, E.M., Villaescusa, J.C., et al. (2016). Molecular Diversity of Midbrain Development in Mouse, Human, and Stem Cells. Cell 167, 566–580.e19. 10.1016/j.cell.2016.09.027.

15. Kamath, T., Abdulraouf, A., Burris, S.J., Langlieb, J., Gazestani, V., Nadaf, N.M., Balderrama, K., Vanderburg, C., and Macosko, E.Z. (2022). Single-cell genomic profiling of human dopamine neurons identifies a population that selectively degenerates in Parkinson’s disease. Nat Neurosci 25, 588–595. 10.1038/s41593-022-01061-1.

16. Aristieta, A., Parker, J.E., Gao, Y.E., Rubin, J.E., and Gittis, A.H. (2024). Dopamine depletion weakens direct pathway modulation of SNr neurons. Neurobiology of Disease 196, 106512. 10.1016/j.nbd.2024.106512.

17. Zeng, H. (2022). What is a cell type and how to define it? Cell 185, 2739–2755. 10.1016/j.cell.2022.06.031.

18. Saunders, A., Macosko, E.Z., Wysoker, A., Goldman, M., Krienen, F.M., de Rivera, H., Bien, E., Baum, M., Bortolin, L., Wang, S., et al. (2018). Molecular Diversity and Specializations among the Cells of the Adult Mouse Brain. Cell 174, 1015–1030. 10.1016/j.cell.2018.07.028.

19. Grillner, S. (2021). Evolution of the vertebrate motor system — from forebrain to spinal cord. Current Opinion in Neurobiology 71, 11–18. 10.1016/j.conb.2021.07.016.

20. Leiras, R., Cregg, J.M., and Kiehn, O. (2022). Brainstem Circuits for Locomotion. Annual Review of Neuroscience 45, 63–85. 10.1146/annurev-neuro-082321-025137.

21. Guo, L., Walker, W.I., Ponvert, N.D., Penix, P.L., and Jaramillo, S. (2018). Stable representation of sounds in the posterior striatum during flexible auditory decisions. Nature Communications 9. 10.1038/s41467-018-03994-3.

22. Gandhi, N.J., and Katnani, H.A. (2011). Motor functions of the superior colliculus. Annual Review of Neuroscience 34, 205–231. 10.1146/annurev-neuro-061010-113728.

23. Cox, J., and Witten, I.B. (2019). Striatal circuits for reward learning and decision-making. Nature Reviews Neuroscience 2019 20:8 20, 482–494. 10.1038/s41583-019-0189-2.

24. Delgado-Zabalza, L., Mallet, N.P., Glangetas, C., Dabee, G., Garret, M., Miguelez, C., and Baufreton, J. (2023). Targeting parvalbumin-expressing neurons in the substantia nigra pars reticulata restores motor function in parkinsonian mice. Cell Reports 42, 113287. 10.1016/j.celrep.2023.113287.

25. Eisinger, R.S., Cernera, S., Gittis, A., Gunduz, A., and Okun, M.S. (2019). A review of basal ganglia circuits and physiology: Application to deep brain stimulation. Parkinsonism & Related Disorders 59, 9–20. 10.1016/j.parkreldis.2019.01.009.

26. Reese, M.G., Eeckman, F.H., Kulp, D., and Haussler, D. (1997). Improved Splice Site Detection in Genie. Journal of Computational Biology 4, 311–323. 10.1089/cmb.1997.4.311.

27. Azim, E., Jiang, J., Alstermark, B., and Jessell, T.M. (2014). Skilled reaching relies on a V2a propriospinal internal copy circuit. Nature 508, 357–363. 10.1038/nature13021.

28. Deverman, B.E., Pravdo, P.L., Simpson, B.P., Kumar, S.R., Chan, K.Y., Banerjee, A., Wu, W.-L., Yang, B., Huber, N., Pasca, S.P., et al. (2016). Cre-dependent selection yields AAV variants for widespread gene transfer to the adult brain. Nat Biotechnol 34, 204–209. 10.1038/nbt.3440.

29. Schindelin, J., Arganda-Carreras, I., Frise, E., Kaynig, V., Longair, M., Pietzsch, T., Preibisch, S., Rueden, C., Saalfeld, S., Schmid, B., et al. (2012). Fiji: an open-source platform for biological-image analysis. Nat Methods 9, 676–682. 10.1038/nmeth.2019.

30. Botta, P., Fushiki, A., Vicente, A.M., Hammond, L.A., Mosberger, A.C., Gerfen, C.R., Peterka, D., and Costa, R.M. (2020). An Amygdala Circuit Mediates Experience-Dependent Momentary Arrests during Exploration. Cell 183, 605–619.e22. 10.1016/j.cell.2020.09.023.

31. Nelson, A., Abdelmesih, B., and Costa, R.M. (2021). Corticospinal populations broadcast complex motor signals to coordinated spinal and striatal circuits. Nature Neuroscience 24. 10.1038/s41593-021-00939-w.

32. Thévenaz, P., Ruttimann, U.E., and Unser, M. (1998). A pyramid approach to subpixel registration based on intensity. IEEE Trans Image Process 7, 27–41. 10.1109/83.650848.

33. Sommer, C., Straehle, C., Kothe, U., and Hamprecht, F.A. (2011). Ilastik: Interactive learning and segmentation toolkit. In 2011 IEEE International Symposium on Biomedical Imaging: From Nano to Macro (IEEE), pp. 230–233. 10.1109/ISBI.2011.5872394.

34. Klein, S., Staring, M., Murphy, K., Viergever, M.A., and Pluim, J.P.W. (2010). elastix: a toolbox for intensity-based medical image registration. IEEE Trans Med Imaging 29, 196–205. 10.1109/TMI.2009.2035616.

35. Ragan, T., Kadiri, L.R., Venkataraju, K.U., Bahlmann, K., Sutin, J., Taranda, J., Arganda-Carreras, I., Kim, Y., Seung, H.S., and Osten, P. (2012). Serial two-photon tomography for automated ex vivo mouse brain imaging. Nat Methods 9, 255–258. 10.1038/nmeth.1854.

36. Chon, U., Vanselow, D.J., Cheng, K.C., and Kim, Y. (2019). Enhanced and unified anatomical labeling for a common mouse brain atlas. Nat Commun 10, 5067. 10.1038/s41467-019-13057-w.

37. Isolation of Nuclei from Adult Mouse Brain Tissue Protocol v1 (2019). 10.17504/protocols.io.7dxhi7n.

38. Stuart, T., Butler, A., Hoffman, P., Hafemeister, C., Papalexi, E., Mauck, W.M., Hao, Y., Stoeckius, M., Smibert, P., and Satija, R. (2019). Comprehensive Integration of Single-Cell Data. Cell 177, 1888–1902.e21. 10.1016/j.cell.2019.05.031.

39. Zappia, L., and Oshlack, A. (2018). Clustering trees: a visualization for evaluating clusterings at multiple resolutions. GigaScience 7. 10.1093/gigascience/giy083.

40. Dobin, A., Davis, C.A., Schlesinger, F., Drenkow, J., Zaleski, C., Jha, S., Batut, P., Chaisson, M., and Gingeras, T.R. (2013). STAR: ultrafast universal RNA-seq aligner. Bioinformatics 29, 15–21. 10.1093/bioinformatics/bts635.

41. Korsunsky, I., Millard, N., Fan, J., Slowikowski, K., Zhang, F., Wei, K., Baglaenko, Y., Brenner, M., Loh, P., and Raychaudhuri, S. (2019). Fast, sensitive and accurate integration of single-cell data with Harmony. Nat Methods 16, 1289–1296. 10.1038/s41592-019-0619-0.

